# Long-term Production and Recovery of Medium-Chain Carboxylates from Source-Separated Organics

**DOI:** 10.64898/2026.03.25.714070

**Authors:** Diana Dyussekenova, Jasmeen K. Parmar, Marjan A. Ezabadi, Blake G. Lindner, Youngseck Hong, Jay R. Werber, Christopher E. Lawson

**Affiliations:** Department of Chemical Engineering & Applied Chemistry, University of Toronto, Toronto, ON, Canada; Veolia Water Technologies, Oakville, ON, Canada

## Abstract

Source-separated organics (SSO) are widely processed via anaerobic digestion to produce biogas, yet alternative conversion pathways could generate higher-value products. Here, we demonstrate long-term continuous production and recovery of medium-chain carboxylic acids (MCCAs) from SSO via microbial chain elongation using a bench-scale anaerobic bioreactor operated for 911 days. The bioreactor was fed with SSO samples collected from two full-scale municipal organics processing facilities in Toronto, Canada, capturing facility-specific and seasonal variability in SSO composition. MCCA production was closely associated with the availability of lactate as an electron donor, which varied with SSO preprocessing operations and outdoor collection temperatures. To mitigate product inhibition, an in-line extraction system using hollow-fiber polydimethylsiloxane (PDMS, also known as silicone polymer) membranes was integrated with the anaerobic membrane bioreactor, providing a robust and solvent-free alternative to solvent-based extraction methods. Selective extraction reduced MCCA product inhibition when lactate availability was not limiting, resulting in maximum MCCA yields and production rates of 0.31 g MCCA g^-1^ VS_feed_ and 0.84 g L^-1^ day^-1^, respectively, with notable octanoic acid production (up to 20% of total MCCA). Acidification of the alkaline extract produced a phase-separated MCCA-rich oil (0.88 g MCCA g^-1^ oil) without the addition of downstream separation steps. Microbial community analysis of the bioreactor revealed enrichment of putative chain-elongating bacteria, including *Eubacterium* and *Pseudoramibacter* species, while shifts in SSO feedstock microbiomes influenced substrate availability and product spectra. These results demonstrate the feasibility of sustained MCCA production from municipal organic waste streams and highlight opportunities to integrate chain elongation with existing anaerobic digestion infrastructure.

## 1. INTRODUCTION

Transforming organic waste into a wide range of valuable resources is a central challenge for achieving resource recovery within a circular bioeconomy. Approximately 20% of global methane emissions are attributed to landfilling and waste management.^1^ In Canada, organic waste landfilling accounts for approximately 17% of annual methane emissions.^2^ Anaerobic digestion (AD) is a common waste diversion strategy used to reduce the environmental impacts of landfilling and recover energy in the form of biogas, which requires separating organics from the general municipal solid waste stream for ease of valorization. Municipal green bin programs, implemented in several countries including Canada, collect food waste and other organic materials, such as plant waste, pet waste and soiled paper, separately from the general municipal solid waste stream.^3^ These programs enable residents to produce a separate stream of organic waste, referred to as source-separated organics (SSO), which can be directly transported to full-scale AD facilities for treatment (e.g., AD facilities in Toronto, Canada process ∼120,000 t SSO yr^-1^). However, the low market value of biogas, even when upgrading to renewable natural gas (∼$0.74 kg^-1^), limits the broader application of AD.^4,5^

Production of medium-chain carboxylic acids (MCCAs) from SSO via microbial chain elongation represents an opportunity to expand the product spectrum of traditional AD facilities beyond biogas towards higher-value chemicals. MCCAs are monocarboxylic acids containing 6 to 12 carbons, including hexanoic acid (C6), heptanoic acid (C7), and octanoic acid (C8). They have garnered interest as a target product for resource recovery due to their relatively high market value (approximately $4 kg^-1^ C6),^4^ potential for upgrading to other high-value oleochemicals,^4,6^ and need to replace existing unsustainable palm-based refinery processes.^7^ In the AD food web, intermediates, including lactate, ethanol, and short-chain carboxylic acids (SCCAs; C2–C5), are produced from the hydrolysis of organic waste, and can be used by chain-elongating bacteria for MCCA production via the reverse β-oxidation (RBO) pathway.^6^ The RBO pathway includes the generation of acetyl-CoA from an electron donor (such as lactate or ethanol), which enters a cyclic process and elongates carboxylic acids with two carbons at a time, making even-chain MCCAs from acetate (C2) and odd-chain MCCAs from propionate (C3).^5,6,8^

Several groups have demonstrated MCCA production from various organic waste streams (e.g., food waste,^9–11^ lignocellulosic residues,^12^ dairy processing waste,^13^ and ethanol-rich waste streams^14,15^) via chain elongation at the bench scale. These feedstocks can differ in their organic composition (e.g., carbohydrates, proteins, lipids, and readily available electron donors), which can influence MCCA yields and product spectrum.^16^ To improve substrate availability and promote chain elongation, pre-treatment strategies and supplementation with external electron donors (e.g., ethanol) are sometimes employed, which can increase operating costs and the environmental footprint of MCCA production.^17^ A few studies have utilized food waste for MCCA production without supplying external electron donors.^9–11^ Some studies have utilized SSO for biogas and SCCA production;^18–20^ however, the impacts of seasonal variations in SSO composition on MCCA production have not been investigated.

Additionally, the accumulation of undissociated MCCAs has been shown to inhibit microbial activity and limit production rates, with reported toxicity thresholds of ∼7.5 mM undissociated C6 at pH 5.5.^6,14^ Consequently, continuous MCCA extraction systems have been employed to maintain MCCA concentrations below inhibitory levels. Various membrane-based in-line extraction technologies have been integrated with MCCA bioreactor systems;^21^ however, solvent-based liquid-liquid extraction (e.g., pertraction) has been the most widely employed approach due to its selectivity for longer-chain carboxylic acids.^14,15,22–26^ Polydimethylsiloxane (PDMS) membranes have recently been shown to exhibit MCCA permeabilities and selectivity of MCCAs over SCCAs comparable to those of solvent-based extraction systems, with estimated operating costs approximately 12-fold lower due to their solvent-free nature and lower membrane area requirements.^27^ However, the application of this technology for long-term MCCA extraction from complex organic waste streams remains largely unexplored. Furthermore, phase separation of MCCAs as an oil product has been demonstrated using electrochemical separation technologies, including electrolysis,^25,28^ and electrodialysis.^22,29^ Phase separation can also be achieved through acidification by exploiting the lower solubilities of MCCAs relative to SCCAs. While acidification may lower capital costs associated with downstream phase separation, it requires additional chemical inputs and generates salt by-products.

In this study, we demonstrate the feasibility of achieving stable MCCA production and recovery from complex SSO waste without the addition of external electron donors in a continuous bench-scale bioreactor system. The bioreactor was operated continuously for 911 days while using SSO samples collected from two full-scale organics processing facilities in Toronto, enabling evaluation of the impacts of seasonal and facility-specific variations in feedstock composition on MCCA production performance. Partway through operation, a solvent-free in-line MCCA extraction system based on PDMS hollow-fiber membranes was integrated to mitigate MCCA inhibition and facilitate downstream recovery. The impacts of SSO feedstock composition and changes in process conditions (solids retention time, in-line extraction) on MCCA titer, production rate, and yield metrics were evaluated. Finally, the microbial community profiles of the feedstock and bioreactor across all process conditions were characterized. Our findings provide important insights into factors governing high MCCA production from SSO and the challenges associated with processing real organic waste streams, informing the design and operation of larger-scale systems.

## 2. MATERIALS AND METHODS

### 2.1. Bioreactor Influent and Inoculum

SSO samples were collected from two full-scale organics processing facilities in Toronto, Dufferin and Disco Road. These facilities utilize AD to treat SSO waste (green-bin waste) collected from households and public buildings, with annual processing capacities of 55,000 t yr^-1^ and 75,000 t yr^-1^, respectively.^20,30^ At the facilities, solid green-bin waste is pre-processed prior to feeding the digesters. This process involves removal of inorganic material (e.g., plastic bags), homogenization with process water, and grit removal, producing a liquid slurry, referred to as SSO pulp, with a total solids content varying between 6–9%. For this study, the SSO pulp produced immediately after pre-processing was collected from the facilities approximately once a month and stored at 4°C. Prior to use in the bioreactor, the SSO pulp was sieved through a 0.5 mm sieve to remove large materials that could clog bioreactor tubing. SSO pulp samples from Dufferin were used during Days 1–441 of bioreactor operation, while SSO pulp samples from Disco Road were used during Days 442–911 of operation. The bioreactor was inoculated once at start-up (Day 1) with a 1:1 (v/v) mixture of Dufferin SSO sample (substrate) and Dufferin suspension buffer tank sample (inoculum). The suspension buffer tank sample was chosen as the inoculum since it contained high MCCA concentrations (details are provided in S1). No additional inoculation was performed during bioreactor operation, including after the transition from Dufferin to Disco Road feedstock. The characteristics of the inoculum and SSO samples throughout the operational periods are provided in Table S1.

### 2.2. Bioreactor System

Continuous fermentation was carried out under anaerobic conditions in a 5 L jacketed bioreactor (Minifors, INFORS HT). The bioreactor was operated at 37°C and a pH of 5 was maintained by automated addition of 1 M sulfuric acid and/or 1 M potassium hydroxide. Sulfate concentrations were monitored during Days 261–447, with no substantial reduction observed in the bioreactor (see further details in S2). Sieved SSO pulp was fed continuously and maintained at approximately 4°C. The feed bottle was replaced periodically and sparged with nitrogen gas prior to feeding, to maintain anaerobic conditions. The bioreactor was sparged with nitrogen gas before start-up, and during process upsets (air intrusion) on Days 89, 201, 252, 772–779. The feed and effluent flow rates were controlled by peristaltic pumps connected to programmable timers to maintain a constant working volume and achieve target hydraulic retention time (HRT) and solids retention time (SRT). Working volume was decreased from 2.86 L to 2.70 L on Day 307 and increased to 3.40 L on Day 488 to allocate space for ultrafiltration (UF) membranes. The bioreactor was operated as a continuous stirred-tank reactor (CSTR) until Day 625, after which continuous membrane filtration was integrated, converting the system into an anaerobic membrane bioreactor (AnMBR).

### 2.3. Extraction System

Submerged UF membranes (Veolia Water Technologies & Solutions, ON, Canada, 0.04 µm pore size, 0.045 m^2^ membrane surface area of each module) were used for solids-liquid separation in the AnMBR, enabling biomass retention and independent control of HRT and SRT, while minimizing fouling of the downstream PDMS membranes. Regular back pulsing with permeate was implemented on Day 659 (every 12 minutes for 30 seconds) as a cleaning strategy in place. PDMS hollow-fiber membranes (PDMSXA, PermSelect, MI, USA, 0.83 m^2^ membrane surface area of each module, 55-µm wall thickness) were used for in-line extraction. AnMBR permeate was passed through the lumen side of the PDMS membranes and recirculated back to the AnMBR (Figure 1). An alkaline solution made up of sodium hydroxide (0.02 mM NaOH, pH ∼9; Sigma-Aldrich, St. Louis, MO, USA) was recirculated at approximately 430 L day^-1^ on the shell side of PMDS membranes to extract MCCAs. Starting on Day 636, the pH of the alkaline solution was automatically controlled at a pH of 9 using 3 M sodium hydroxide with a dosing pump.

**Figure 1.**
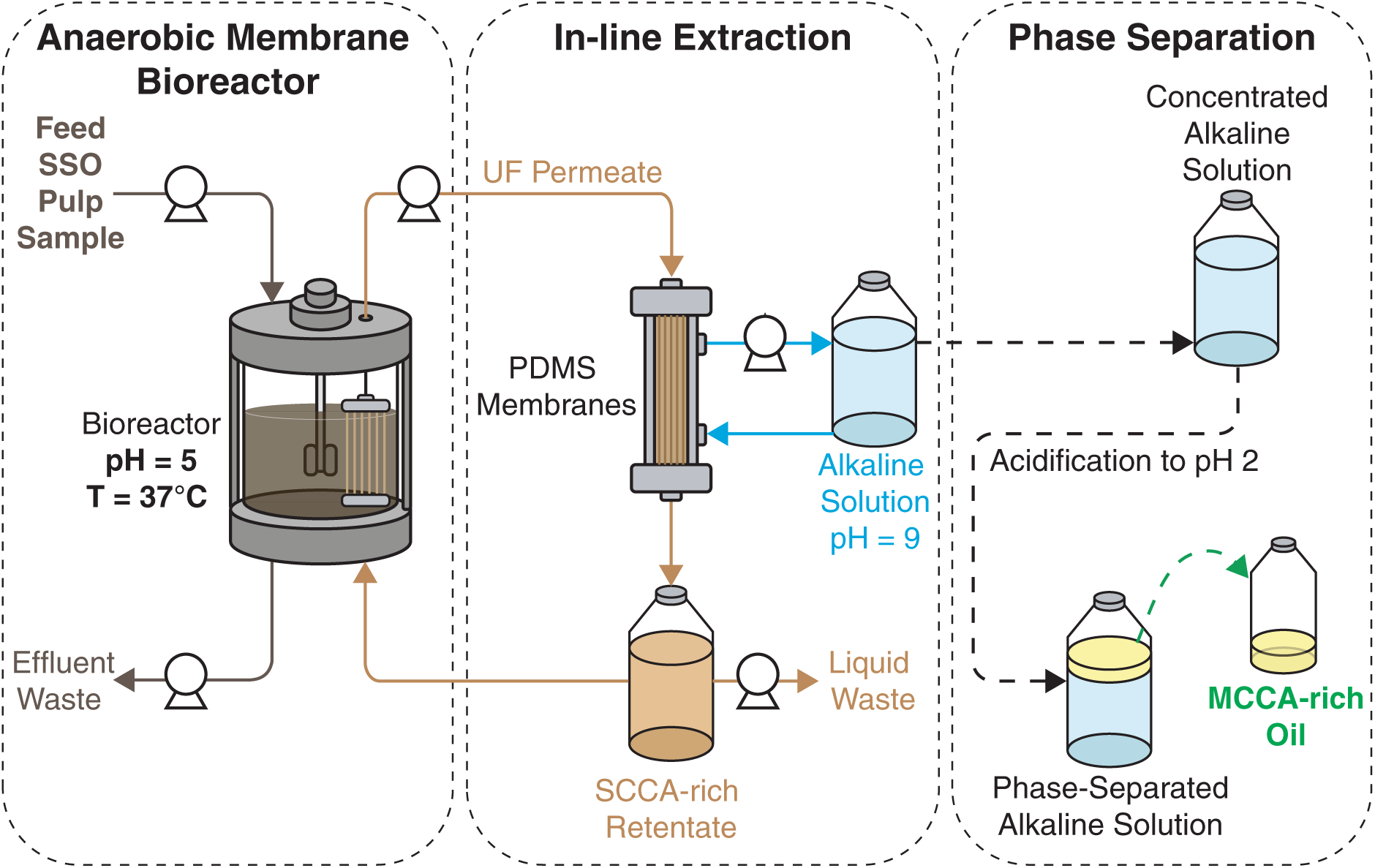
Schematic representation of the continuous bioreactor with submerged ultrafiltration membranes (AnMBR) integrated with an in-line extraction system and MCCA phase separation. Phase separation was done in batch mode.

The in-line extraction was integrated with the bioreactor and operated from Days 557–563, Days 574–582 and from Days 625–911 with some interruptions from technical difficulties (detailed information is provided in S3). On Day 663, PDMS membrane area was increased from 1.66 m^2^ to 2.49 m^2^ and the UF membrane area was increased from 0.045 m^2^ to 0.09 m^2^ to improve the extraction efficiency. On Day 784, two additional hollow-fiber PDMS membrane modules were added in series (M60-B, Nagasep, Sumida, Japan, 0.98 m^2^ total membrane surface area, 60-µm wall thickness).

Acidification of the alkaline solution for MCCA-oil recovery was performed by the addition of 6 M hydrochloric acid (Sigma-Aldrich) to a final pH of 2 (Figure 1). Detailed information on acidification is provided in S4. Following phase separation, organic and aqueous layers were characterized by carboxylic acid concentrations and chemical oxygen demand (COD).

### 2.4. Operating Conditions

The bioreactor operating time was divided into eight periods reflecting the changes in operating conditions, SSO source and feedstock outdoor collection temperatures (Table 1). An initial start-up period occurred during Days 1–55, operating at ∼7-day SRT and HRT. Period 1 (Days 56–146) corresponded to 8-day SRT and HRT, Periods 2a–3b (Days 147–624) corresponded to 12-day SRT and HRT, Periods 4a and 4b (Days 625–811) corresponded to 12-day SRT and HRT with in-line extraction, and Period 5 (Days 812–911) corresponded to 25-day SRT and 12-day HRT with in-line extraction. In Periods 1–2b, SSO samples were sourced from Dufferin. Starting from Day 442, SSO samples were sourced from Disco Road and used in Periods 3a–5. Periods 1, 2b, 3a, 4b, and 5 represent outdoor collection temperature of SSO feedstock above 16°C (referred to as warm), while Periods 2a, 3b and 4a represent outdoor collection temperature below 13°C (referred to as cold). Average performance metrics presented in Table 1 were calculated across pseudo steady-state operation, defined as 3 times HRT, specifically when there was a change in SRT or SSO source (Periods 1, 2a, 3a and 5). Values are reported as mean ± standard deviation and represent temporal variations within each operating period. Detailed calculations of performance metrics are provided in S5.

**Table 1.**
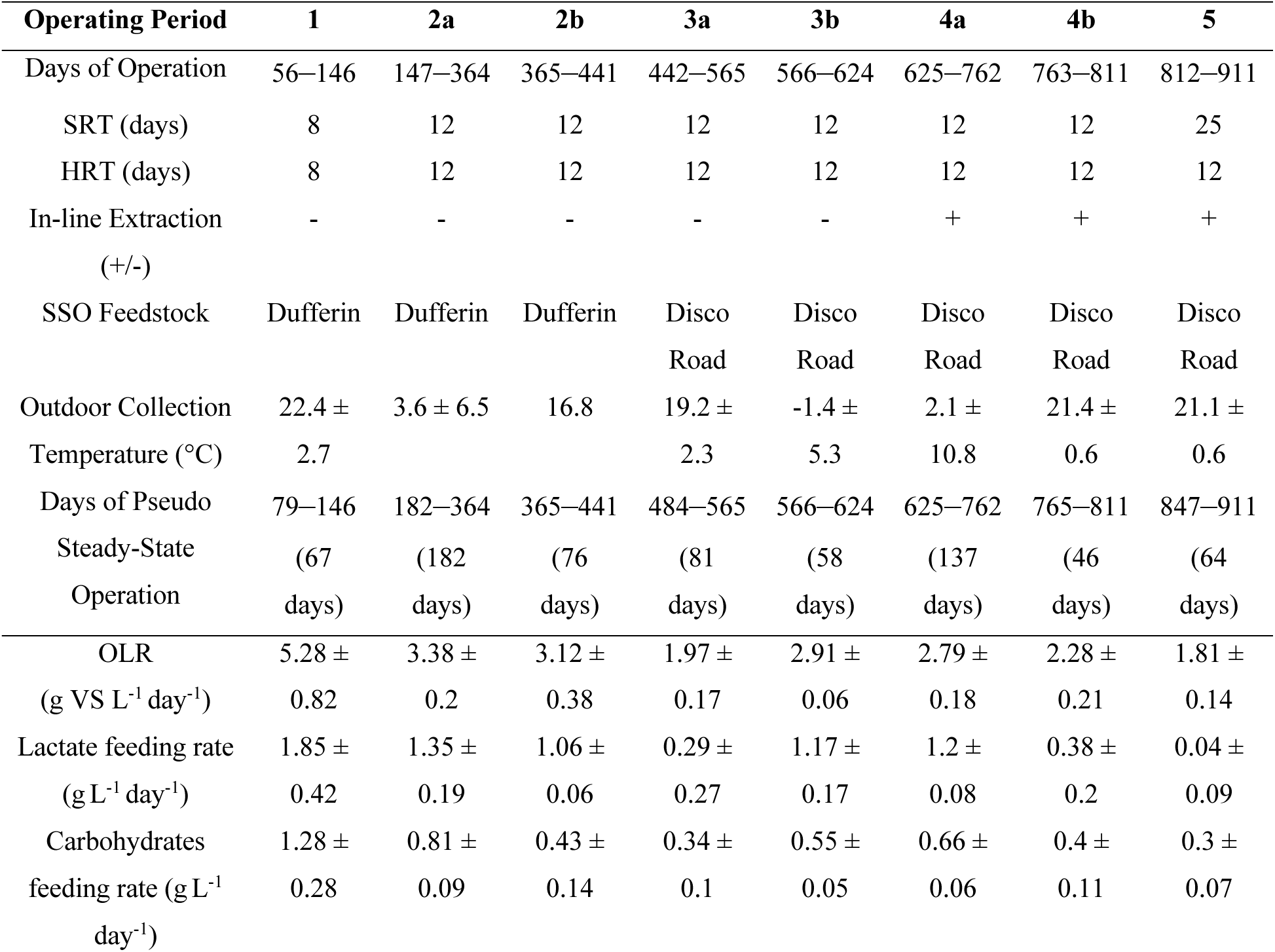
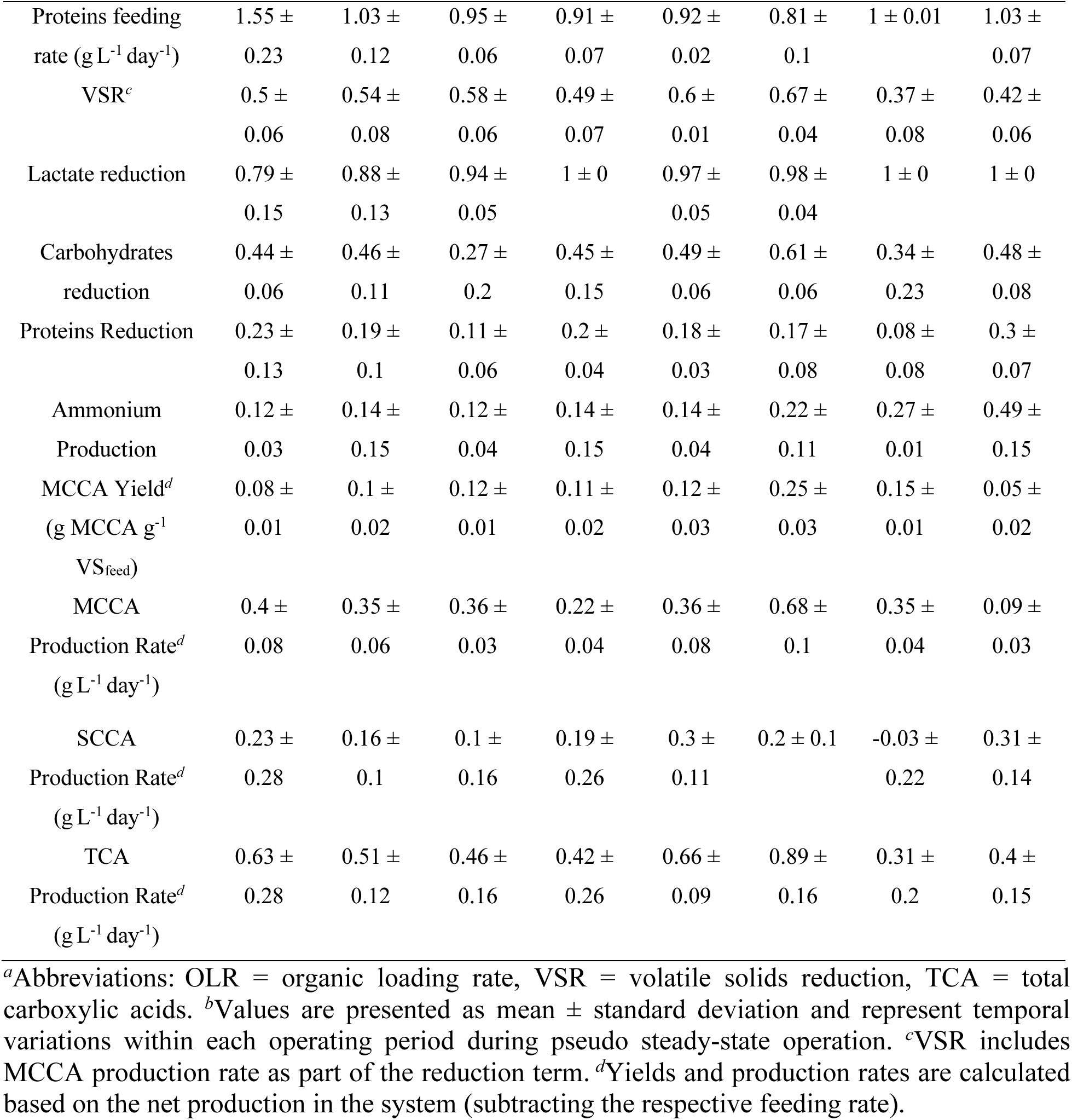
Summary of experimental periods, operating conditions, and performance metrics*^a,b a^*Abbreviations: OLR = organic loading rate, VSR = volatile solids reduction, TCA = total carboxylic acids. *^b^*Values are presented as mean ± standard deviation and represent temporal variations within each operating period during pseudo steady-state operation. *^c^*VSR includes MCCA production rate as part of the reduction term. *^d^*Yields and production rates are calculated based on the net production in the system (subtracting the respective feeding rate).

### 2.5. Chemical Analyses

Bioreactor and feedstock samples were collected daily, centrifuged (10,000 rcf for 10 minutes) and supernatant was filtered through 0.22 µm for chromatographic analyses. Biomass was stored at −20°C for 16S rRNA gene sequencing. Permeate after PDMS membranes and alkaline solution were sampled daily after the measurement of permeate flow rate (UF flux). Carboxylic acids and ethanol were quantified using Agilent 8890 gas chromatograph (GC) equipped with tandem mass spectrometry (7000D Triple Quadrupole, Agilent, Santa Clara, CA, USA), and lactate was measured using high-performance liquid chromatography (UltiMate 3000 HPLC System, Thermo Scientific, Waltham, MA, USA) with details provided in S6. Ammonium was measured using ion chromatography (IC) and colorimetric quantification assay (data consistency between two methods is provided in S7). Bioreactor headspace composition (CO_2_, CH_4_, O_2_, N_2_) was analyzed weekly on GC (Hewlett Packard 5890, Agilent) with a thermal conductivity detector (TCD) and CTR I Concentric Packed Column (Alltech, Nicholasville, KY, USA) using helium carrier gas, while H_2_ was verified on Days 208, 210, 310, 324, 337, 373 and 462 (detailed information in S8). Chemical and solids analyses were done once a week. Solids (TS, VS, TSS, VSS) were determined according to Standard Methods.^31^ The soluble portion of the samples was filtered through 0.45 µm after centrifugation. Soluble COD (sCOD) and total COD (tCOD) were measured using COD digestion vials (Method 8000, Hach, Loveland, CO, USA). Soluble and total carbohydrates were quantified using the phenol-sulfuric method with glucose standard. Soluble and total proteins were measured using Pierce BCA protein assay kit (Thermo Scientific).

### 2.6. Microbial Community Analysis

SSO feedstock samples collected from Dufferin and Disco Road across eight operating periods were selected for full-length bacterial (n = 8) and archaeal (n = 7, Period 1 samples did not amplify) 16S rRNA gene sequencing. Corresponding bioreactor effluent samples collected during operation with the respective SSO feedstocks were also included. For bacterial 16S rRNA gene sequencing, two bioreactor effluent samples were selected from each operating period (n = 16), with samples separated by no more than three SRTs. For archaeal 16S rRNA gene sequencing, one bioreactor effluent sample was selected from each operating period except Period 1 (n = 7).

DNA extraction was performed using DNeasy PowerSoil Pro Kit (Qiagen, Hilden, Germany) according to the manufacturer’s protocol. Extracted DNA samples were quantified with Qubit dsDNA HS Assay Kit using Qubit 4 fluorometer (Invitrogen, Waltham, MA, USA). PCR amplification was performed using normalized DNA extracts (∼40 ng template input), 2X Platinum SuperFi II Green PCR Master Mix (Invitrogen), forward and reverse primers, in a 50 μL reaction volume and consisting of 25 cycles. Bacterial 16S rRNA gene primers included 27F (AGRGTTYGATYMTGGCTCAG) and 1391R (GACGGGCGGTGWGTRCA),^32–34^ archaeal primers included SSU1ArF (TCCGGTTGATCCYGCBRG) and SSU1000ArR (GGCCATGCAMYWCCTCTC).^33,35^ Amplification was performed twice, and duplicate samples were pooled for clean-up using Monarch PCR & DNA Cleanup Kit (New England Biolabs, Ipswich, MA, USA). Cleaned products were quantified using Quant-iT dsDNA BR Assay kit (Invitrogen). Nanopore sequencing libraries of samples and ZymoBIOMICS Microbial Community DNA Standard (Zymo Research, Irvine, CA, USA) were prepared using ONT Native Barcoding Kit 96 V14 (SQK-NBD114.96) according to the manufacturer’s instructions and sequenced on the MinION R10.4.1 flow cell (Oxford, United Kingdom) with separate libraries prepared and sequenced for bacterial and archaeal 16S amplicons. Detailed information on bioinformatics and statistical analyses is provided in S9.

## 3. RESULTS

### 3.1. Impact of seasonal variations in SSO composition on MCCA production

SSO samples collected from two organics processing facilities, Dufferin and Disco Road, over 911 days of bioreactor operation provided a unique opportunity to investigate the effect of seasonal and facility-specific variability on SSO waste characteristics, and the subsequent impact on MCCA production. Seasonal effects on SSO were found to be primarily temperature-driven and were represented by the changes in average outdoor temperatures during the feedstock collection days.

SSO pulp samples from Dufferin were used on Days 1–441 (Periods 1–2b), spanning both 8-day and 12-day SRT operations. In Periods 1 and 2b, Dufferin SSO pulp samples were collected during warm outdoor temperatures, 22.4 ± 2.7°Cand 16.8°C, respectively, while in Period 2a, during cold outdoor temperatures, 3.6 ± 6.5°C. Across these periods, the chemical composition and preservation of SSO pulp samples were not substantially impacted by the outdoor collection temperature, represented by high lactate (16.22 ± 3.9 g COD L^-1^) and low SCCA concentrations (6.71 ± 2.58 g COD L^-1^) (Figure 2). During Periods 2a and 2b, the bioreactor was operated at a 12-day SRT, allowing the comparison of MCCA performance metrics with respect to the chemical composition of Dufferin feedstock. MCCA production rates in Period 2a and 2b were similar, 0.35 ± 0.06 g L^-1^ day^-1^ and 0.36 ± 0.03 g L^-1^ day^-1^ (Table 1), respectively, with C6 as the dominant MCCA produced (Figure 3A). Lactate was primarily consumed, while ethanol reduction was not observed (Figure S10), suggesting that the availability of lactate in the Dufferin feedstock contributed to the stable MCCA production observed in these periods.

**Figure 2.**
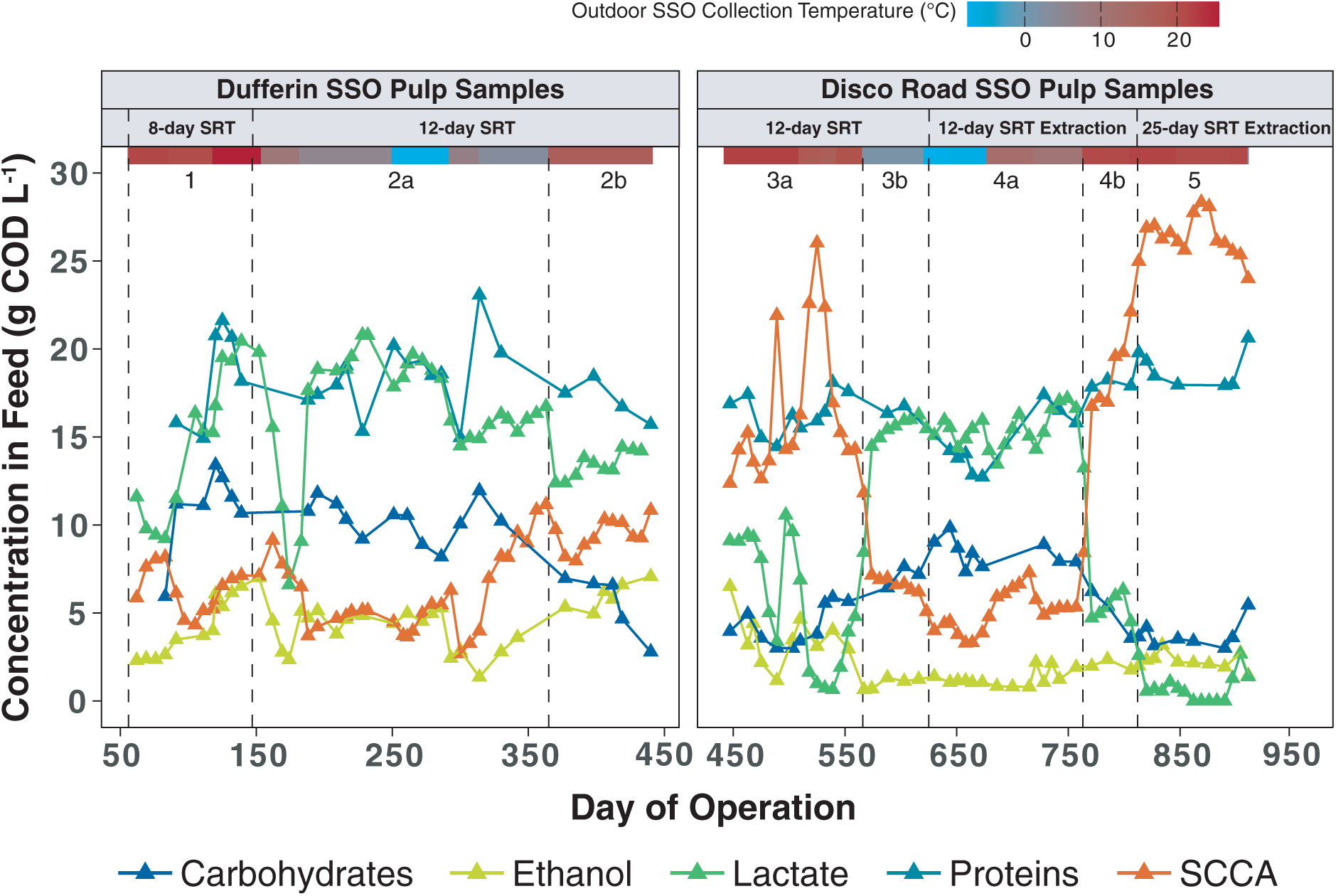
Chemical characterization profiles of SSO pulp samples from Dufferin and Disco Road organics processing facilities (total carbohydrates in blue, ethanol in light green, lactate in green, total proteins in teal and SCCAs in orange). Operating periods are separated by dashed vertical lines. Outdoor collection temperature of SSO samples is presented as temperature gradient (blue represented colder periods and red represents warmer periods). Sample points for lactate, ethanol and SCCAs include 5-day averages, while carbohydrates and proteins were not averaged as characterized not daily.

**Figure 3.**
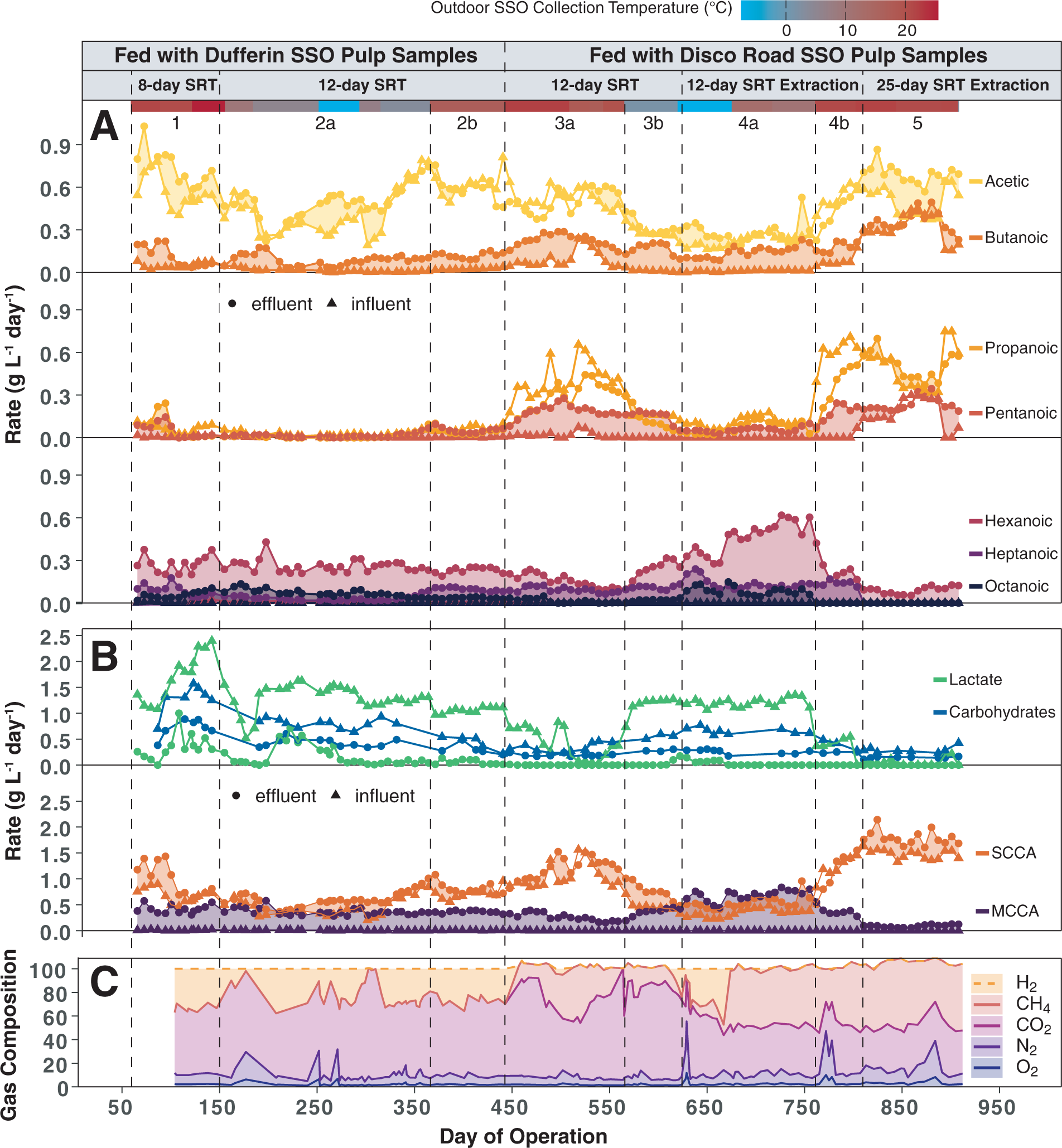
Bioreactor performance over operating periods (dashed vertical lines): (A) feeding (triangles) and effluent (circles) rates of carboxylic acids, with filled regions representing production rates, (B) feeding (triangles) and effluent (circles) rates of lactate, carbohydrates, SCCAs and MCCAs, (C) bioreactor headspace composition (hydrogen was estimated from headspace balance and verified on selected days; nitrogen was normalized on days when it was introduced to the system, detailed information is provided in S7). Carboxylic acids and lactate sample points are 5-day averages, while carbohydrates are not averaged as characterized not daily. Effluent rates from Period 4a include extraction rates.

SSO pulp samples from Disco Road were used on Days 442–911 (Periods 3a–5), spanning the 12-day SRT operation, addition of in-line extraction, and 25-day SRT operation. Periods 3a (19.2 ± 2.3°C), 4b (21.4 ± 0.6°C) and 5 (21.1 ± 0.6°C) represent Disco SSO samples collected during warm periods, while Periods 3b (−1.4 ± 5.3°C) and 4a (2.1 ± 10.8°C) represent cold periods. In contrast to Dufferin SSO samples, the chemical composition of Disco feedstock varied with outdoor collection temperatures. Concentrations of lactate and carbohydrates were lower during warm periods than during cold periods (Figure 2). During warm periods, lactate and carbohydrate concentrations averaged 3.51 ± 3.35 g COD L^-1^ and 4.19 ± 1.08 g COD L^-1^, respectively, compared with 15.31 ± 1.32 g COD L^-1^ and 8.08 ± 0.94 g COD L^-1^ during cold periods. On the other hand, SCCA concentrations were higher during warm periods (20.56 ± 5.8 g COD L^-1^) than during cold periods (5.48 ± 1.33 g COD L^-1^) (Figure 2), with C3 dominating the feedstock carboxylic acid spectrum (Figure S10). These results suggest that Disco SSO samples were more susceptible to degradation past lactate during warmer periods than Dufferin pulp.

Variations in Disco feedstock composition observed between warm and cold collection periods were associated with differences in the MCCA product spectrum and production rates. During Periods 3a and 3b, the bioreactor was operated at a 12-day SRT receiving warm and cold Disco feedstock, respectively. In Period 3a (warm), the lactate feeding rate was lower and the C3 feeding rate was higher than in Period 3b (cold) (Figure 3A,B). Correspondingly, the C6 production rate was lower in Period 3a (0.13 ± 0.03 g L^-1^ day^-1^) than in Period 3b (0.23 ± 0.07 g L^-1^ day^-1^), while C7 production rates were similar (Figure 3A). In Period 3b, the lactate consumption rate (1.21 ± 0.18 g COD L^-1^ day^-1^) was substantially higher than the ethanol consumption rate (0.08 ± 0.03 g COD L^-1^ day^-1^) (Figure S10), suggesting that the availability of lactate as an electron donor contributed to the increased C6 production observed in Period 3b. Additionally, the comparable C7 production rates observed in Periods 3a and 3b, despite lower lactate availability in Period 3a, suggest that the elevated C3 feeding rate during the warm period potentially promoted odd-chain elongation.

### 3.2. Impact of switching feedstock source from Dufferin to Disco Road

SSO feedstock source was switched from Dufferin (Period 2b) to Disco (Period 3a) on Day 442 during warm outdoor collection temperatures, enabling comparison of bioreactor performance under feedstocks originating from the two facilities. This change reduced the OLR from 3.12 ± 0.38 to 1.97 ± 0.17 g VS_feed_ L^-1^ day^-1^ (Table 1). In Period 3a, the loading rates of lactate and carbohydrates decreased, and the C3 loading increased (Figure 3A,B). Correspondingly, odd-chain carboxylate product formation increased, including an increase in C5 production from 0.05 ± 0.02 to 0.18 ± 0.05 g L^-1^ day^-1^ (Figure 3A) and an increase in the C7 fraction of total MCCA production from 27 ± 3% to 39 ± 2%. MCCA production rate decreased from 0.36 ± 0.03 g L^-1^ day^-1^ to 0.22 ± 0.04 g L^-1^ day^-1^ (Table 1). Dufferin pulp samples maintained higher concentrations of lactate than Disco samples at comparably warm outdoor temperatures. These results indicate that the Dufferin pulp samples were preferable for achieving higher MCCA production, since they were more stable against degradation of carbohydrates beyond lactate at varying outdoor temperatures. A potential factor causing these differences may be the SSO collection and distribution system at the facilities. For example, longer residence time during green bin collection (e.g., due to longer transit time between transfer stations) could accelerate the degradation of SSO prior to its arrival at the tipping floor of the processing facility.

Gas composition in the bioreactor headspace also differed between operation with Dufferin and Disco Road SSO pulp. Hydrogen was detected during periods with Dufferin feedstock at partial pressures of ∼0.25 atm (Figure 3C). Previous studies have suggested that hydrogen accumulation is not expected to directly limit MCCA production, although its partial pressure may influence metabolic pathways utilized by the microbial community.^36^ After the transition from Dufferin to Disco feedstock (Day 442), methane was first observed in the bioreactor headspace on Day 457 and continued to be detected for the remainder of the study, occupying a substantial fraction of the headspace gas composition (Figure 3C). The emergence of methane suggests differences in archaeal methanogenic potential between the SSO feedstocks, particularly in Disco samples. This is potentially due to differences in facility practices. For example, methanogens in Disco SSO samples may have originated from recycling of untreated digester effluent streams (e.g., process water from digester centrate, or digester solids) back into the SSO pulpers.

### 3.3. Increasing SRT from 8 to 12 days improved MCCA yield with Dufferin feedstock

Next, we evaluated the impact of increasing SRT from 8 to 12 days on MCCA production. To minimize the effect of seasonal variability in the feedstock composition and subsequent bioreactor performance, we compared metrics from Periods 1 (8-day SRT) and 2b (12-day SRT), which occurred using Dufferin feedstock collected during comparable warm outdoor temperatures. During Period 1 (8-day SRT), MCCA yield averaged 0.08 ± 0.01 g MCCA g^-1^ VS_feed_, with incomplete consumption of the available lactate from the feed (79 ± 15%) (Table 1). MCCA yield and lactate consumption increased during Period 2b (12-day SRT), reaching 0.12 ± 0.01 g MCCA g^-1^ VS_feed_ and 94 ± 5%, respectively, despite operating at a lower OLR (Table 1). These results suggest that increasing SRT to 12 days while using Dufferin feedstock, which maintained stable lactate concentrations, improved MCCA yield. This improvement may be attributed to longer retention of slower-growing chain-elongating bacteria, enabling improved lactate utilization during this operational period.

MCCA production rates were comparable in Periods 1 and 2b, 0.4 ± 0.08 g L^-1^ day^-1^ and 0.36 ± 0.03 g L^-1^ day^-1^, respectively (Table 1). The maximum C6 titer in Period 2b reached 3.35 g L-1 (Day 401), which translates to an undissociated C6 concentration of 12.5 mM (pH of 5), exceeding the previously reported toxicity limit (7.5 mM C6, pH of 5.5).^14^ Therefore, further improvements to the MCCA production rate at 12-day SRT were likely limited due to the toxicity effects of undissociated MCCAs in the bioreactor. This motivated the subsequent addition of continuous in-line MCCA extraction downstream of the bioreactor to reduce the concentration of MCCAs to levels which can alleviate their toxic effects.

### 3.4. Robust extraction efficiency was needed to reduce the toxicity of MCCAs in the bioreactor

This study presents the first demonstration of using hollow-fiber PDMS membranes for in-line MCCA extraction from a real organic waste feedstock. Existing studies often use membrane-based extraction (pertraction) for MCCA separation,^14,15,22–26^ where a hydrophobic solvent (usually mineral oil) with or without an organic extractant (e.g., TOPO) is used to sorb undissociated MCCAs, and an alkaline solution is used to desorb MCCAs as carboxylate salts. In contrast, solvents and extractants are not required for extraction with PDMS membranes, as undissociated MCCAs diffuse through the membrane layer and desorb into the alkaline solution. Previously, our team showed that PDMS membranes have promising permeability selectivities of longer-chain relative to shorter-chain carboxylic acids (e.g., ∼233 for C8/C2), which are comparable to solvent-based extraction.^27^

In-line extraction was implemented on Day 625 (Period 4a) to reduce MCCA toxicity in the bioreactor and demonstrate the recovery of MCCAs in the oil phase. Due to high solids concentration in the bioreactor (14.28 ± 1.51 g TSS L^-1^, Days 447–623), submerged UF membranes were used to produce solids-free permeate (i.e., AnMBR) and prevent fouling of the downstream PDMS membranes. The permeate stream was continuously recirculated on the lumen side of the PDMS membranes, while an alkaline solution (pH of 9) was used on the shell side to drive MCCA extraction (Figure 1).

Reducing MCCA concentration in the AnMBR relied on robust extraction efficiency, defined as the ratio of MCCA extraction rate to MCCA production rate. This overall extraction efficiency depended on both the permeate flow rate (UF flux) and the efficiency of PDMS membranes to selectively extract MCCAs. Here, PDMS efficiency was defined as the ratio of the MCCA extraction rate to the MCCA loading rate entering the membrane modules with the permeate stream and depended on membrane area, MCCA/SCCA selectivities, and MCCA permeabilities. Initially (Days 625–663), MCCA extraction efficiency was low (0.34 ± 0.11 g g^-1^), resulting in no observed reduction in the MCCA titer in the AnMBR (4.42 ± 0.57 g L^-1^, Figure 4A,B) compared with Period 3b prior to extraction implementation (4.23 ± 1 g L^-1^). PDMS efficiency showed moderate MCCA separation (0.47 ± 0.1 g g^-1^) (Figure 4A), while UF flux remained low (0.3 ± 0.1 LMH), limiting the rate of extraction. Therefore, further optimization of the extraction efficiency was needed to reduce MCCA titers below toxicity levels.

**Figure 4.**
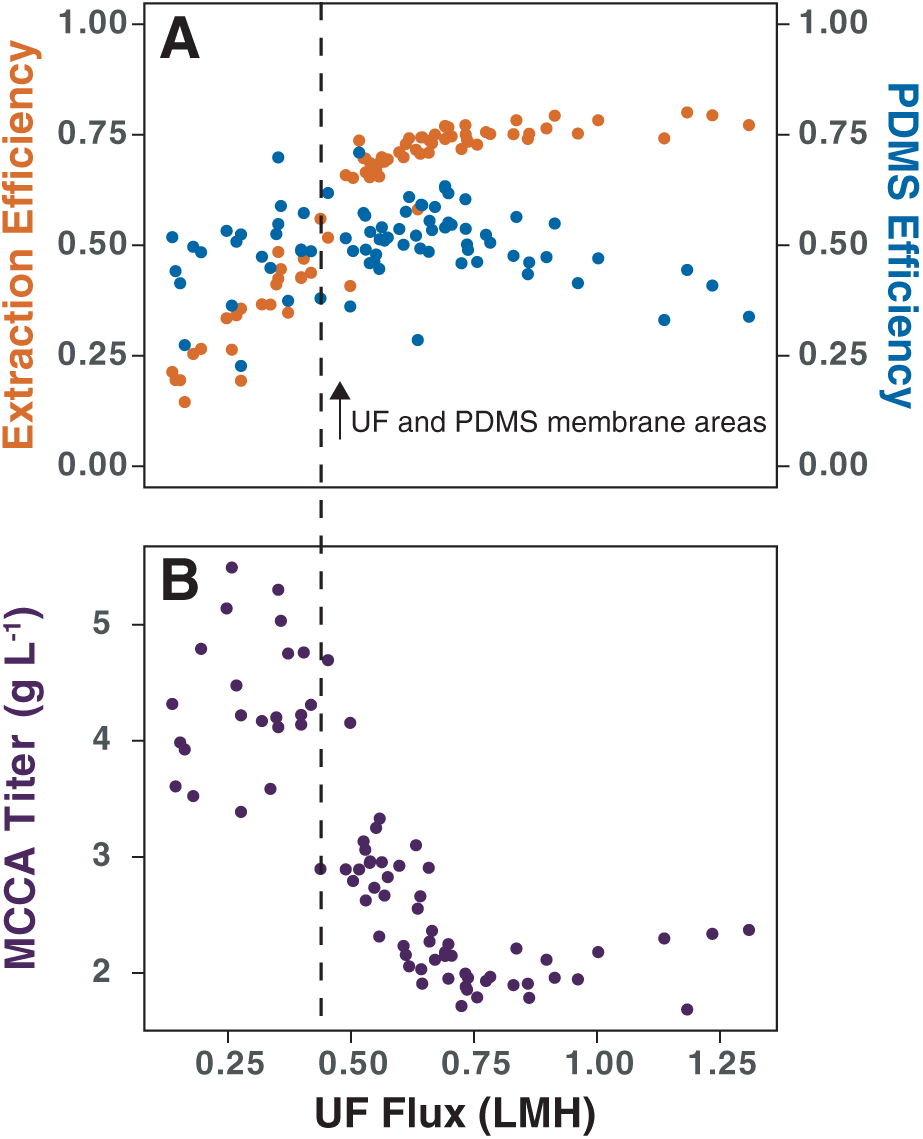
Extraction performance as a function of UF flux (Period 4a, Days 625–762): (A) extraction efficiency (orange) as g MCCA extracted over g MCCA produced and PDMS efficiency (blue) as g MCCA extracted over g MCCA introduced to membranes, and (B) MCCA titer in the AnMBR (purple).

Optimization on Day 663 included increasing both UF and PDMS membrane areas by 2-fold and 1.5-fold, respectively. The goal was to increase permeate flow rate (UF flux) while maintaining the residence time within PDMS membranes despite faster permeate loading. As a result, UF flux was improved by approximately two-fold (0.69 ± 0.18 LMH, Days 665–762) and PDMS efficiency was also maintained (0.51 ± 0.08 g g^-1^) (Figure 4A). These changes resulted in improved extraction efficiency (0.73 ± 0.04 g g^-1^) and approximately two-fold decrease in MCCA titer in the AnMBR (2.36 ± 0.45 g L^-1^) (Figure 4A,B).

### 3.5. Continuous MCCA extraction increased MCCA yield and productivity

The AnMBR integrated with in-line extraction receiving cold Disco SSO pulp operated from Day 625–762 (Period 4a). Prior to extraction implementation, after three SRTs during Period 3b (Days 609–623), MCCA yield and production rate remained stable at 0.15 g MCCA g^-1^ VS_feed_ and 0.43 ± 0.01 g L^-1^ day^-1^, respectively, providing a baseline in MCCA production performance for comparison. Additionally, OLR and lactate feeding rate in Period 4a were similar to Period 3b (Table 1), allowing the comparison of performance metrics with respect to in-line extraction implementation under comparable feedstock composition.

MCCA yield and production rate increased during the first two weeks of in-line extraction implementation, reaching 0.28 g MCCA g^-1^ VS_feed_ and 0.76 g L^-1^ day^-1^ on Day 639, respectively. As the extraction efficiency dropped afterwards, due to a decrease in UF membrane performance, MCCA yield and production rate also decreased, and on average were 0.22 ± 0.04 g MCCA g^-1^ VS_feed_ and 0.59 ± 0.12 g L^-1^ day^-1^, respectively (Days 625–663). To improve MCCA extraction efficiency, additional PDMS and UF membrane areas were added on Day 663 (1.5-fold and 2-fold increase in membrane area, respectively). This resulted in a maximum MCCA yield of 0.31 g MCCA g^-1^ VS_feed_ on Day 673 and a maximum MCCA production rate of 0.84 g L^-1^ day^-1^ on Day 728. The improved MCCA production performance was consistent with previous studies employing in-line extraction, where enhanced MCCA production has been attributed to reduced product inhibition.^14,15,24^ MCCA yield and production rate averaged 0.26 ± 0.02 g MCCA g^-1^ VS_feed_ and 0.72 ± 0.06 g L^-1^ day^-1^, respectively, for the remainder of Period 4a (Days 665–762). The drop in performance was due to fouling of the UF membranes, leading to an uncontrolled decrease in extraction efficiency by approximately 0.1 g g^-1^ and a subsequent increase in MCCA titer by approximately 0.9 g L^-1^ from Day 665 to 762. If the extraction efficiency was controlled, the MCCA yield would likely be maintained close to the maximum achieved in this period.

During Period 4a (with extraction, cold Disco pulp), the production rate of even-chain MCCAs improved, compared to Period 3b (no extraction, cold Disco pulp). Maximum C6 and C8 productivities were achieved, reaching 0.62 g L^-1^ day^-1^ and 0.15 g L^-1^ day^-1^, respectively (Figure 3A). On average, C6 production rate improved from 0.23 ± 0.07 to 0.46 ± 0.11 g L^-1^ day^-1^, while C8 improved from 0.02 ± 0.01 to 0.09 ± 0.03 g L^-1^ day^-1^ compared to Period 3b. C7 production rates remained similar between Periods 3b and 4a (0.11 ± 0.01 and 0.14 ± 0.04 g L⁻¹ day⁻¹, respectively), while the C5 production rate decreased from 0.15 ± 0.05 to 0.06 ± 0.02 g L⁻¹ day⁻¹. These results suggest that in-line extraction promoted selectivity towards longer even-chain carboxylate products when electron donors, such as lactate, were preserved in the SSO pulp.

Production rate of total carboxylic acids (TCA) increased from 0.66 ± 0.09 to 0.89 ± 0.16 g L^-1^ day^-1^ in Period 4a compared to Period 3b, with more selectivity toward MCCA production (0.68 ± 0.1 g L^-1^ day^-1^) (Table 1). Lactate and protein reduction did not change, while carbohydrates reduction slightly increased, from 0.49 ± 0.06 to 0.61 ± 0.06 g g^-1^ (Table 1). As carbohydrates hydrolysis was likely improved, more readily available electron donors, such as soluble sugars, were potentially released for lactic acid and chain-elongating bacteria. Additionally, recycling non-extracted SCCAs back to the AnMBR provided more time for their further elongation as described in previous studies,^15^ potentially contributing to increased MCCA selectivity in Period 4a, when electron donors remained available in the feedstock.

### 3.6. Feedstock composition during warmer periods limited MCCA production despite in-line extraction

As the feedstock transitioned from cold to warm Disco pulp (Period 4b, Days 763–811), AnMBR performance with in-line extraction became comparable to Period 3a (no extraction, warm Disco pulp). Similarly to Period 3a, during Period 4b, the lactate feeding rate decreased while the C3 feeding rate increased, resulting in an increased C5 production, reaching 0.18 ± 0.05 g L^-1^ day^-1^ (Figure 3A,B). Production rates of C6 decreased to 0.21 ± 0.04 g L^-1^ day^-1^, while C7 remained relatively unchanged, and C8 was not produced during Period 4b. MCCA yield and production rate decreased to 0.15 ± 0.01 g MCCA g^-1^ VS_feed_ and 0.35 ± 0.04 g L^-1^ day^-1^, respectively (Table 1). As Period 4b operated for a shorter duration (46 days) than Period 3a (81 days), the higher MCCA production metrics observed in Period 4b may partially reflect the shorter operational duration. Additionally, despite a further increase in PDMS membrane area on Day 784, MCCA production rate did not increase, indicating that substrate availability, rather than extraction performance, was likely the primary limitation during Period 4b.

The SRT was increased to 25 days (Period 5, Days 811–911) to evaluate whether longer biomass retention could improve the hydrolysis of residual organic matter in Disco SSO during warmer periods. However, despite the increased SRT and continued in-line extraction, operation with warm-period Disco feedstock, characterized by reduced lactate availability and higher SCCA concentrations (Figure 2), resulted in elevated SCCA production, mainly C2 and C3, and further decreased MCCA production (Figure 3A; see further discussion in S11).

### 3.7. High-purity MCCA oil recovered from alkaline bottle acidification without additional downstream process units

Extraction of undissociated MCCAs from the AnMBR was driven by a concentration gradient across the PDMS membrane, which was maintained by the pH gradient between the AnMBR (pH 5) and the alkaline solution (pH 9). The alkaline pH promoted dissociation of the extracted MCCAs into their corresponding carboxylate ions. Carboxylate salts are highly soluble in water, allowing them to accumulate at concentrations that exceed the solubility limits of the corresponding protonated acids (10.82 g L^-1^ C6, 2.4 g L^-1^ C7, 0.68 g L^-1^ C8).^6^ This accumulation facilitates downstream phase separation, which occurs when the concentrated MCCA salts are protonated by lowering the pH below the pKa values (4.88 C6 and C7, 4.89 C8).^6^

MCCAs were selectively concentrated in the alkaline solution, reaching 0.87 ± 0.02 g MCCA g^-1^ TCA (Days 730–771), predominantly as C6 (Figure 5A). This alkaline solution was used to demonstrate the recovery of an MCCA oil product. MCCA concentrations in the alkaline bottle prior to acidification were 17.84 g L^-1^ C6, 4.75 g L^-1^ C7, and 4.61 g L^-1^ C8 (Day 771). Following acidification to a pH of 2 and phase separation, MCCAs were predominantly recovered in the oil phase (0.65 g MCCA_oil_ g^-1^ TCA_total_), resulting in 86% MCCA recovery (Figure 5B). In contrast, SCCAs remained predominantly in the aqueous phase after acidification (0.05 g SCCA_oil_ g^-1^ TCA_total_) due to the substantially higher aqueous solubilities of their protonated forms compared with MCCAs. MCCA recovery in the oil phase could potentially be improved by concentrating the alkaline bottle for a longer duration, allowing higher concentrations of MCCA carboxylate salts to accumulate prior to acidification. For example, to achieve 95% recovery, the estimated MCCA concentration would need to exceed 200 g L^-1^ (calculations are provided in S4). The phase-separated oil in this study was highly pure, reaching 0.88 g MCCA g^-1^ oil, and was mostly composed of extracted carboxylic acids (Figure 5C). Overall, these results demonstrate that MCCA-rich oil can be recovered through simple acidification, with comparable purities to those in studies that used electrochemical separation.^25,29^

**Figure 5.**
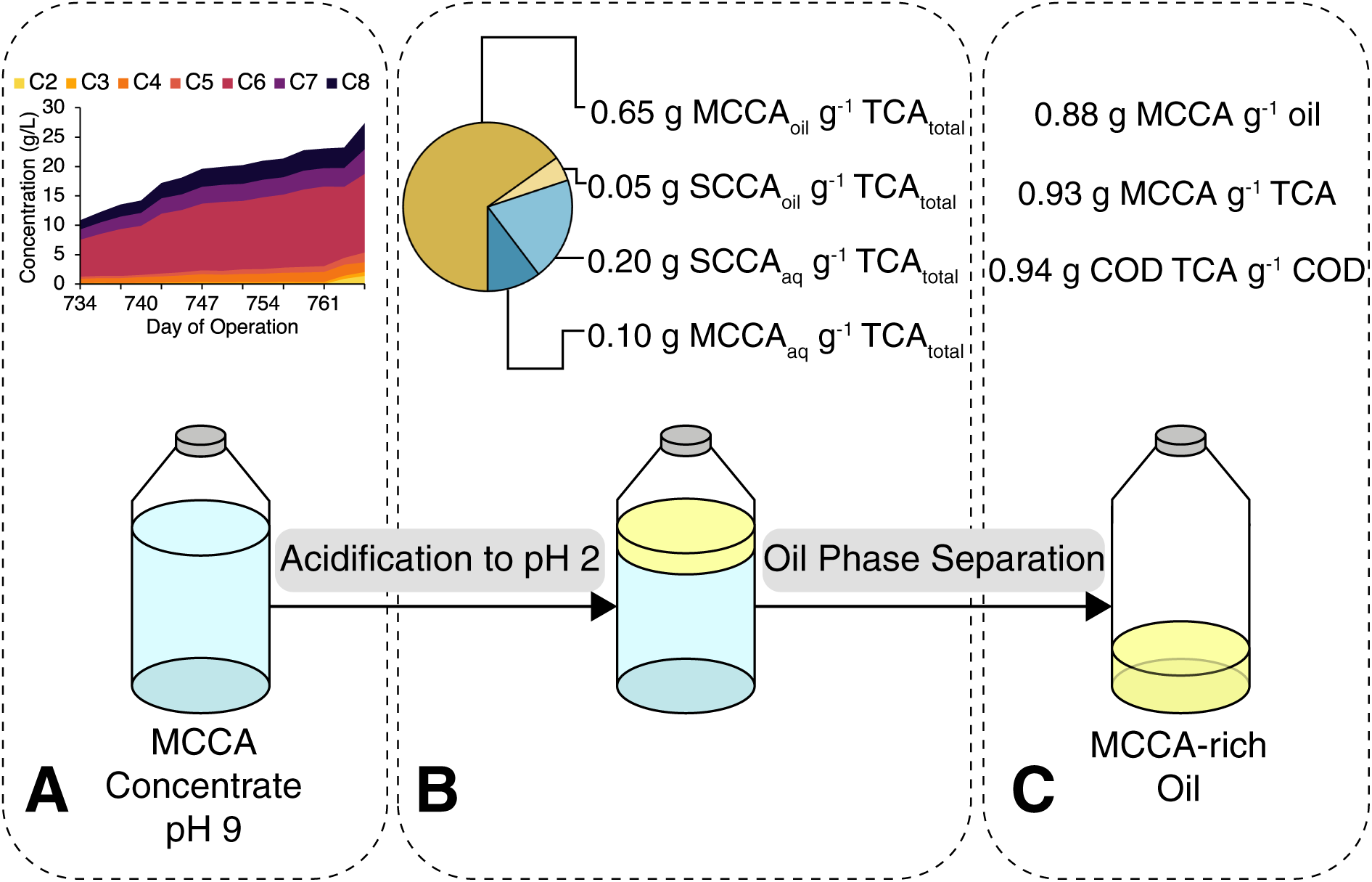
MCCA-oil recovery: (A) selective extraction of MCCAs over SCCAs in the alkaline bottle (Days 730–771), (B) distribution of MCCAs and SCCAs post-acidification per total carboxylic acids (TCA) in oil and aqueous phases and (C) characterization of phase-separated oil.

### 3.8. Microbial composition in the feedstock correlates with observed variations in chemical composition

Microbial community composition of the Dufferin SSO samples remained relatively stable over time and across different outdoor collection temperatures, whereas variations were observed in the Disco samples. The Dufferin microbial community was primarily dominated by *Lactobacillus* (Figure 6A), a well-described genus of lactic acid bacteria (LAB),^37^ consistent with stable high lactate concentrations observed in the feedstock. In contrast, the microbial community profile of Disco SSO samples varied across collection periods (Figure 6A), coinciding with observed fluctuations in feedstock chemical composition. Among LAB genera, *Streptococcus*,^37^ was dominant when lactate was preserved (cold Disco, Days 566 and 719). However, during warmer periods, putative SCCA-producing populations were detected and LAB relative abundances decreased. On Days 442 and 763 (warm Disco), *Prevotella,* a genus containing putative propionate-producing populations,^38,39^ was present at low relative abundance in the Disco SSO. During this time, C3 concentrations in SSO samples increased while lactate concentrations decreased, suggesting that propionate-producing populations may have been active despite their low relative abundance. The Day 847 sample (warm Disco) was separated from the remaining Disco feedstock samples in the NMDS ordination (Figure S12). During this time, *Megasphaera* was relatively abundant alongside *Prevotella.* Of the reads assigned to *Megasphaera*, 89.3% were classified as *M. elsdenii,* a species associated with C3 production when lactate is available and C4 and C5 production when lactate is depleted.^40,41^ Additionally, *Lachnospiraceae NK3A20 group,* which has been previously associated with C4 production when grown on glucose,^42^ was detected at low relative abundance. During this time, SCCA concentrations in Disco SSO were the highest and lactate was fully degraded. Archaeal communities also differed between facilities. In Disco SSO samples, *Methanosaeta,* a genus associated with acetoclastic methanogenesis,^43–45^ dominated across all time points (Figure 6B). In contrast, *Candidatus Nitrocosmicus*, a putative ammonia-oxidizing archaeon,^46^ was dominant in Dufferin samples. Overall, microbial community composition differed between SSO samples from the two facilities (PERMANOVA, R² = 0.53, p = 0.011, Table S13), consistent with the observed differences in feedstock chemical composition and further supporting that facility-specific processing conditions influenced feedstock quality.

**Figure 6.**
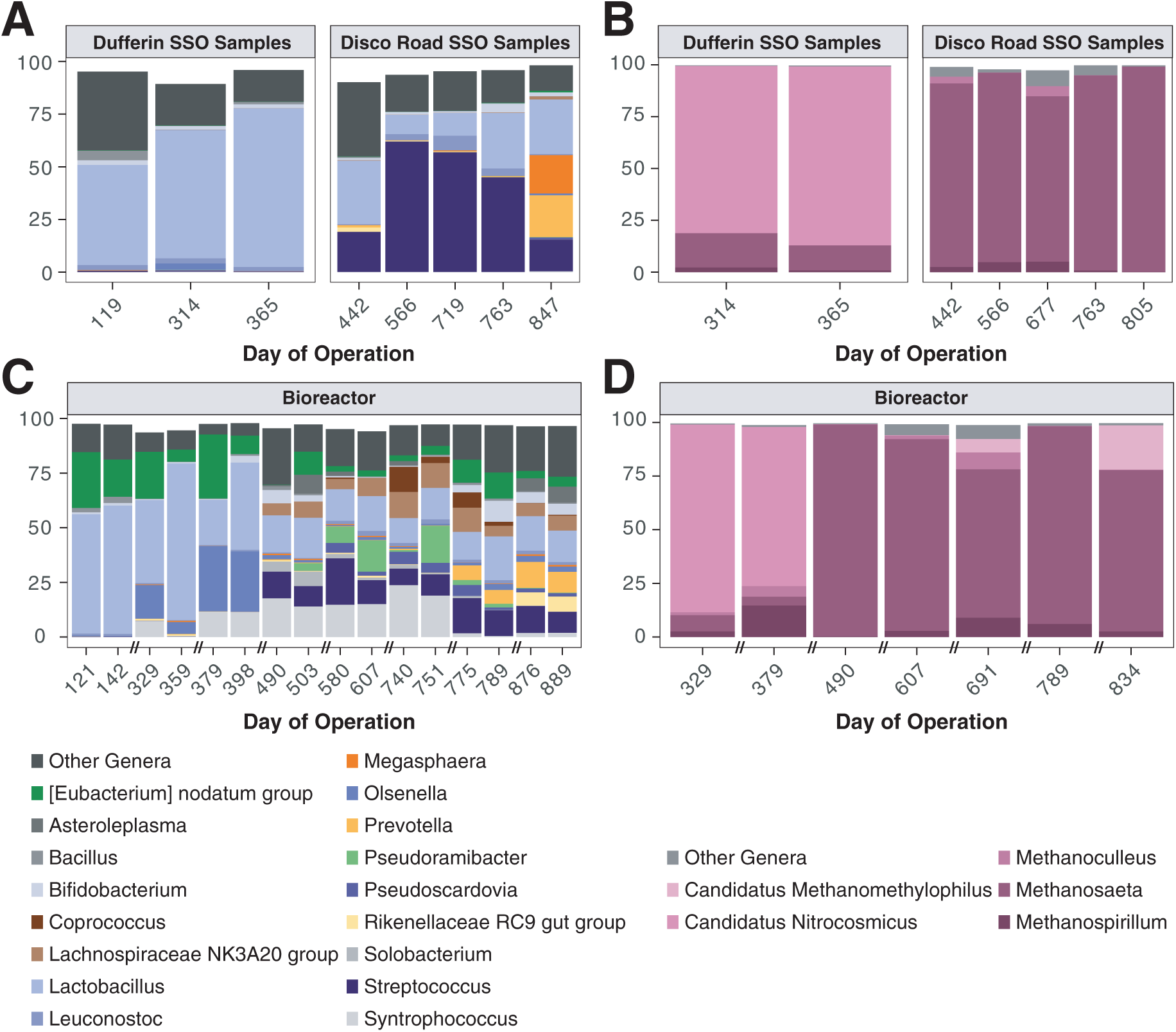
Relative abundance as % of top 17 bacterial groups and top 5 archaeal groups classified to the genus level: (A) bacterial composition in SSO feed samples, (B) archaeal composition in SSO feed samples, (C) bacterial composition in the bioreactor and (D) archaeal composition in the bioreactor. The residual relative abundance belongs to groups not classified down to the genus level.

### 3.9. Bioreactor microbial community profile correlates with observed fluctuations in MCCA production

Microbial community profile in the bioreactor were primarily dominated by the genera *Lactobacillus, Eubacterium*, and *Olsenella* during operation with Dufferin SSO (Days 121–398) (Figure 6C). At the species level, 100% of reads assigned to genus *Eubacterium* were classified as the unnamed species *Eubacterium_T sp903789635,* which was previously binned from metagenomic sequencing of a chain-elongating reactor microbiome.^47^ Belonging to the same genus is the isolated species *E. pyruvativorans,* which has been previously reported as chain-elongating bacteria (CEB).^36,48^ Following the transition to Disco SSO, bioreactor community diversity increased (Shannon, Figure S14), coinciding with the emergence of additional bacterial taxa. *Pseudoramibacter*, with >90% of reads within this genus classified as *Pseudoramibacter sp022484225*, was detected in the bioreactor microbial profile (Days 503, 580, 607, 751), and *Eubacterium* persisted but at lower abundance. *P. alactolyticus,* belonging to the same genus as *P. sp022484225,*^47^ has been reported as MCCA-producing CEB that can utilize lactate.^8,36^

MCCA production fluctuated during operation with Disco SSO samples and coincided with variations in feedstock quality. Specifically, MCCA production decreased in Period 4b and reached the lowest values during Period 5. Starting from Day 775 (Period 4b), the relative abundance of *Prevotella* increased in the bioreactor (Figure 6C). Additionally, on Days 876 and 889 (Period 5), *Rikenellaceae RC9 gut group* also became more abundant. Both *Prevotella* and the *Rikenellaceae RC9 gut group* contain populations of putative SCCA producers, with the latter uncultivated group associated with polysaccharide degradation and acetate production in rumen.^38,39,49^ During Period 5, acetate was the dominant carboxylic acid produced, and propionate production was observed in the bioreactor (Figure 3A). *Lachnospiraceae NK3A20,* a putative group of butyrate-producing organisms,^42^ was present across all operating periods with Disco SSO samples, consistent with greater C4 production observed in the bioreactor. Lastly, the archaeal community was dominated by the putative acetoclastic methanogens (Figure 6D), *Methanosaeta*,^43–45^ supporting the observed methane production throughout operation with Disco SSO and indicating that methanogenesis was not completely inhibited at low pH.

Overall, bioreactor microbial community clustered according to the feedstock source facility (Figure S12), with facility explaining the largest proportion of variation in community composition among the factors examined (PERMANOVA, R² = 0.53, p = 0.002, Table S13). These results further support the importance of feedstock characteristics in shaping bioreactor microbial community structure.

## 4. DISCUSSION

### 4.1. MCCA production rate improved with in-line extraction and was dependent on the availability of lactate in the feedstock

Existing studies have reported increased MCCA production with in-line extraction, attributed to the reduced toxicity from undissociated MCCAs.^14,15,24^ Before in-line extraction was introduced (Period 3b, cold Disco pulp), the maximum C6 titer reached 4.10 g L^-1^ (35.31 mM) on Day 607, which translates to the undissociated concentration of 15.2 mM at a pH of 5. This is higher than the previously reported toxicity limit of 7.5 mM C6 at a pH of 5.5,^14^ suggesting that MCCA production prior to extraction implementation in this study was likely influenced by product inhibition. MCCA production rate with extraction was increased to 0.73 ± 0.04 g L^-1^ day^-1^ (5.97 ± 0.84 mmol L^-1^ day^-1^, 1.62 ± 0.24 g COD L^-1^ day^-1^, Days 665–762). To achieve this productivity, the estimated undissociated MCCA concentration in the bioreactor would need to be ∼31 mM (assuming no extraction), exceeding previously reported toxicity thresholds and suggesting that reducing MCCA titers facilitated improved chain elongation.

During the extraction period in this study (Period 4a, cold Disco pulp), OLR was 5.45 ± 0.44 g COD L^-1^ day^-1^, from which lactate accounted for 1.28 ± 0.09 g COD L^-1^ day^-1^ (Figure S10). Based on complete lactate consumption and a stoichiometric ratio of 3 mol lactate to 1 mol C6,^36^ the estimated theoretical C6 production rate was 1.12 ± 0.1 g COD L^-1^ day^-1^, closely matching the measured C6 production, 1.02 ± 0.24 g COD L^-1^ day^-1^ (Table S15). Although ethanol may have also contributed to chain elongation, its consumption rate was substantially lower (0.09 g COD L⁻¹ day⁻¹; Figure S10), suggesting that lactate was the predominant electron donor during this period. In contrast, prior to in-line extraction implementation (Period 3b, cold Disco pulp), estimated theoretical C6 production from observed lactate consumption (1.08 ± 0.16 g COD L^-1^ day^-1^) exceeded the measured production rate (0.51 ± 0.16 g COD L^-1^ day^-1^) while ethanol consumption remained low (0.08 g COD L⁻¹ day⁻¹). Together, these observations suggest that in-line extraction improved selectivity towards MCCA production from available electron donors.

Although the MCCA production rate achieved in this study was lower than previously reported, MCCA production from a complex SSO waste was demonstrated with notable production of C8, up to 20% of total MCCA produced (Period 4a), which is rarely observed in chain-elongation reactor studies. Studies that showed higher MCCA production rates used feedstocks with more readily available electron donors than SSO. *Ge et al.*^14^ reported a C6 production rate of 7.52 ± 0.94 g COD L^-1^ day^-1^ under extraction efficiencies >90% and an OLR of 10.7 g COD L^-1^ day^-1^ using yeast-fermentation beer. Similar performance was observed by *Kucek et al.*^24^ using a synthetic feedstock with lactate and butyrate (C6: 6.9 g COD L^-1^ day^-1^, OLR: 9.1 g COD L^-1^ day^-1^). *Shrestha et al.*^15^ reported improved MCCA production with extraction reaching 13.6 ± 0.42 mmol L^-1^ day^-1^ (∼4.5% C8 based on mmol of carbon produced) using ethanol-rich waste beer and SCCA-rich permeate (OLR: 8.2 g sCOD L^-1^ day^-1^). *Grootscholten et al.*^50^ showed MCCA production of 28.1 g L^-1^ day^-1^ (∼3% of C8 over total MCCA produced) from the organic fraction of municipal solid waste with ethanol supplementation at a short HRT. Reducing HRT while maintaining SRT suitable for hydrolysis provides an opportunity to increase the MCCA productivity if the yield is maintained close to the maximum achieved in this study (0.31 g MCCA g^-1^ VS_feed_). However, this strategy depends on feedstock quality, as during warmer periods with Disco pulp, substrates were converted to SCCAs, reducing lactate availability and subsequently decreasing MCCA selectivity.

### 4.2. MCCA production becomes more limited by the available substrate loading rate than the extraction rate during warmer periods

During warmer periods, MCCA production with Disco SSO feedstock appeared to be limited primarily by substrate availability rather than extraction performance. MCCA production rates during operation with warm Disco feedstock with and without extraction were 0.35 ± 0.04 g L^-1^ day^-1^ (46 days, Period 4b) and 0.22 ± 0.04 g L^-1^ day^-1^ (81 days, Period 3a), respectively, with the former rate slightly higher, likely due to a shorter operation of Period 4b. The estimated undissociated C6 titer in Period 4b under a no-extraction scenario (9.4 mM) remained below the toxicity threshold observed in this study (15.2 mM), suggesting that MCCA inhibition was unlikely to be a major limiting factor.

Instead, warm Disco feedstock appeared to provide limited electron donor availability for chain elongation. During warm periods (Periods 3a, 4b and 5), SCCAs accounted for a major fraction of the OLR (33 ± 6% on a COD basis), while lactate represented only ∼5% of the OLR (Figure S10). Increased SCCA concentrations in the warm Disco feedstock coincided with the detection of *Prevotella* and *Megasphaera*, genera that contain populations associated with SCCA production.^38–41^ As readily available electron donors such as lactate were degraded in warm Disco SSO prior to bioreactor feeding, MCCA production was likely substrate-limited regardless of extraction. Additionally, estimated C6 production from available lactate during operation with warm Disco feedstock (Periods 3a and 4b) aligned with the observed C6 production rates (Table S15), while during operation with Dufferin pulp (Periods 1, 2a, 2b), the observed C6 production rates remained lower than stoichiometric estimates. Together, these observations suggest MCCA production during warm periods with Disco feedstock was limited by lactate availability, while with Dufferin pulp, MCCA production was potentially influenced by product inhibition.

SSO samples sourced from Dufferin appeared less susceptible to substrate degradation beyond lactate, resulting in more stable MCCA production. One of the factors that potentially contributed to these facility-specific variations is the transit time of SSO waste. Overall, these findings highlight that minimizing feedstock degradation is critical for maintaining stable MCCA production, while integration of the extraction system can further enhance production rates and selectivity when sufficient electron donors remain available in the feedstock.

### 4.3. PDMS membranes provide alternative extraction strategy with comparable performance to pertraction

In this study, selective MCCA extraction was demonstrated using PDMS hollow-fiber membranes from the AnMBR treating a real organic waste stream. The MCCA flux averaged 0.72 ± 0.12 g m^-2^ day^-1^ during best performance (Day 665–762) and was comparable to values reported in previous studies employing pertraction. Fluxes from these studies were estimated from reported MCCA production rates and membrane areas for systems achieving extraction efficiencies greater than 90%. Estimated C6 fluxes in studies by *Ge et al.*^14^ and *Kucek et al.*^24^ were 1.0 and 0.7 g m^-2^ day^-1^, respectively. *Zhu et al.*^27^ evaluated a laboratory-made PDMS membrane, with modeling results suggesting performance superior to conventional pertraction systems and lower projected operating costs due to its solvent-free nature and reduced membrane area requirements. Additionally, the authors compared the fluxes of commercial *PermSelect* PDMS membranes (also used in this study) and found that laboratory-made membranes outperformed the commercial membranes by ∼200-fold.^27^ The authors suggested that PDMS fluxes could potentially be improved by tailoring membrane properties, such as material composition, thickness and support layer design,^27^ particularly when considering larger-scale applications. Considering the extraction efficiency achieved in this study was ∼0.73 g g^-1^, it can be potentially improved by 1) increasing MCCA flux as described above 2) increasing the permeate recirculation flowrate though PDMS membrane 3) recycling the bioreactor effluent stream back to the feed line to minimize losses of residual MCCAs leaving the system. However, the second optimization may not be required if higher MCCA fluxes are achieved, as comparable extraction performance could then be maintained at a lower UF flux, which is particularly relevant for systems treating high-solids SSO waste.

### 4.4. Outlook Considerations

In this study, MCCA production and recovery were demonstrated from a complex organic waste stream, providing a foundation for future implementation of this technology as part of product expansion in conventional anaerobic digestion systems. Based on the findings, several optimization strategies will be necessary to achieve high MCCA production rates. First, substrate degradation beyond lactate should be minimized in the SSO feedstock to maintain MCCA/SCCA selectivity. This includes identifying sources of variability in samples from organics processing facilities and evaluating alternative feedstock sources where substrate degradation can be minimized. Second, increasing OLR while maintaining a suitable SRT for hydrolysis may further enhance MCCA production rates. Third, MCCA extraction efficiency can be further improved by optimizing PDMS membrane performance and implementing side-stream recycling to avoid the loss of residual MCCAs in the bioreactor. Maintaining robust extraction efficiency is particularly important for keeping MCCA titer below toxicity limits and ensuring efficient recovery during downstream MCCA oil phase separation. Lastly, future work should include a techno-economic assessment to evaluate the feasibility and scalability of the process.

## Data Availability Statement

16S rRNA gene sequencing data was uploaded to the National Center for Biotechnology Information (NCBI) under BioProject accession number PRJNA1482536.

## Supporting Information

The Supporting Information is available free of charge: Characterization of inoculum and feedstock, preliminary extraction trials and MCCA oil-phase separation, performance metric calculations, carboxylic acid and lactate quantification, ammonium data, gas composition normalization, bioinformatics and statistical analyses of 16S rRNA gene sequencing data, discussion of operation during the 25-day SRT period, and stoichiometric and COD balance data.

## Author Contributions

The manuscript was written through contributions of all authors. C.E.L. and J.R.W. conceptualized the work. D.D and J.K.P led bioreactor operation, experimental analysis, quantification and manuscript writing. M.A.E assisted with bioreactor operation and experimental analysis. B.L assisted with sequencing workflow and led bioinformatics data analysis. Y.H. assisted with conceptualizing the work.

## Acknowledgments

This research was financially supported by Natural Sciences and Engineering Research Council of Canada [ALLRP 580897-22, RGPIN-2021-02684, NSERC-CREATE 528163-2019] through the Alliance Program, Veolia Water Technologies & Solutions (WTS), Envera, CBS Bio Platforms, The Ontario Clean Water Agency (OCWA), and the Ontario Water Consortium (OWC). We thank the City of Toronto for providing SSO access at the Dufferin and Disco Road facilities. We thank Gareth Stock and Nelson Fonseca at Veolia WTS for advice and support. We thank Adriana D’Amico and Enyonam Sewordor for their support in the SSO collection. We thank Joelle Weir, Mara Jezernik, Hazel Fricska, and Yue (Evelyn) Tan for support on bioreactor analyses. We also thank Renzo L. Gutierrez, Jiahao Zhu, Faisal Shahbaz and Hongchen Wang for support with membrane systems.

## For Table of Contents Only

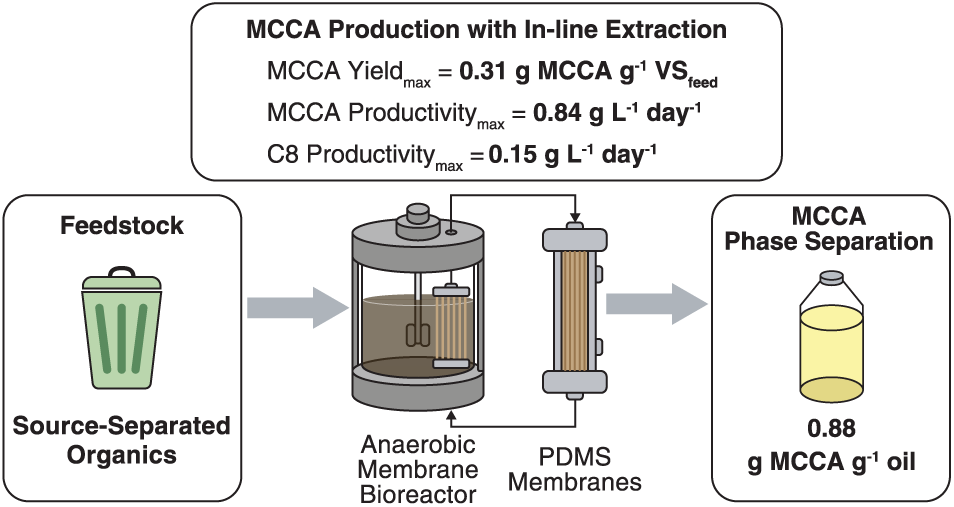

## References

(1) United Nations Environment Programme; Climate & Clean Air Coalition. Global Methane Assessment: Benefits and Costs of Mitigating Methane Emissions; Technical report 978-92-807-3854–4; 2021. https://wedocs.unep.org/handle/20.500.11822/35913.

(2) National Inventory Report 1990-2023: Greenhouse Gas Sources and Sinks in Canada; National Inventory Report; Environment and Climate Change Canada. https://www.canada.ca/en/environment-climate-change/services/climate-change/greenhouse-gas-emissions/inventory.html.

(3) Levis, J. W.; Barlaz, M. A.; Themelis, N. J.; Ulloa, P. Assessment of the State of Food Waste Treatment in the United States and Canada. Waste Management 2010, 30 (8), 1486–1494. 10.1016/j.wasman.2010.01.031.

(4) Scarborough, M. J.; Lawson, C. E.; DeCola, A. C.; Gois, I. M. Microbiomes for Sustainable Biomanufacturing. Current Opinion in Microbiology 2022, 65, 8–14. 10.1016/j.mib.2021.09.015.

(5) Spirito, C. M.; Richter, H.; Rabaey, K.; Stams, A. J.; Angenent, L. T. Chain Elongation in Anaerobic Reactor Microbiomes to Recover Resources from Waste. Current Opinion in Biotechnology 2014, 27, 115–122. 10.1016/j.copbio.2014.01.003.

(6) Angenent, L. T.; Richter, H.; Buckel, W.; Spirito, C. M.; Steinbusch, K. J. J.; Plugge, C. M.; Strik, D. P. B. T. B.; Grootscholten, T. I. M.; Buisman, C. J. N.; Hamelers, H. V. M. Chain Elongation with Reactor Microbiomes: Open-Culture Biotechnology To Produce Biochemicals. Environ. Sci. Technol. 2016, 50 (6), 2796–2810. 10.1021/acs.est.5b04847.

(7) Meijaard, E.; Brooks, T. M.; Carlson, K. M.; Slade, E. M.; Garcia-Ulloa, J.; Gaveau, D. L. A.; Lee, J. S. H.; Santika, T.; Juffe-Bignoli, D.; Struebig, M. J.; Wich, S. A.; Ancrenaz, M.; Koh, L. P.; Zamira, N.; Abrams, J. F.; Prins, H. H. T.; Sendashonga, C. N.; Murdiyarso, D.; Furumo, P. R.; Macfarlane, N.; Hoffmann, R.; Persio, M.; Descals, A.; Szantoi, Z.; Sheil, D. The Environmental Impacts of Palm Oil in Context. Nat. Plants 2020, 6 (12), 1418–1426. 10.1038/s41477-020-00813-w.

(8) Gois, I.; Bowers, C.; Kim, B.-C.; Flick, R.; Lawson, C. Distinct Acetate Utilization Strategies Differentiate Butyrate and Octanoate Producing Chain-Elongating Bacteria; 2025. 10.1101/2025.05.02.651941.

(9) Tang, J.; Yang, H.; Pu, Y.; Hu, Y.; Huang, J.; Jin, N.; He, X.; Wang, X. C. Caproic Acid Production from Food Waste Using Indigenous Microbiota: Performance and Mechanisms. Bioresource Technology 2023, 387, 129687. 10.1016/j.biortech.2023.129687.

(10) Zhang, Y.; Bai, J.; Zuo, J. Performance and Mechanisms of Medium-Chain Fatty Acid Production by Anaerobic Fermentation of Food Waste without External Electron Donors. Bioresource Technology 2023, 374, 128735. 10.1016/j.biortech.2023.128735.

(11) Nzeteu, C. O.; Trego, A. C.; Abram, F.; O’Flaherty, V. Reproducible, High-Yielding, Biological Caproate Production from Food Waste Using a Single-Phase Anaerobic Reactor System. Biotechnology for Biofuels 2018, 11 (1), 108. 10.1186/s13068-018-1101-4.

(12) Scarborough, M. J.; Lynch, G.; Dickson, M.; McGee, M.; Donohue, T. J.; Noguera, D. R. Increasing the Economic Value of Lignocellulosic Stillage through Medium-Chain Fatty Acid Production. Biotechnol Biofuels 2018, 11 (1), 200. 10.1186/s13068-018-1193-x.

(13) Xu, J.; Hao, J.; Guzman, J. J. L.; Spirito, C. M.; Harroff, L. A.; Angenent, L. T. Temperature-Phased Conversion of Acid Whey Waste Into Medium-Chain Carboxylic Acids via Lactic Acid: No External e-Donor. Joule 2018, 2 (2), 280–295. 10.1016/j.joule.2017.11.008.

(14) Ge, S.; Usack, J. G.; Spirito, C. M.; Angenent, L. T. Long-Term n-Caproic Acid Production from Yeast-Fermentation Beer in an Anaerobic Bioreactor with Continuous Product Extraction. Environ. Sci. Technol. 2015, 49 (13), 8012–8021. 10.1021/acs.est.5b00238.

(15) Shrestha, S.; Xue, S.; Kitt, D.; Song, H.; Truyers, C.; Muermans, M.; Smets, I.; Raskin, L. Anaerobic Dynamic Membrane Bioreactor Development to Facilitate Organic Waste Conversion to Medium-Chain Carboxylic Acids and Their Downstream Recovery. ACS EST Eng. 2022, 2 (2), 169–180. 10.1021/acsestengg.1c00273.

(16) Stamatopoulou, P.; Malkowski, J.; Conrado, L.; Brown, K.; Scarborough, M. Fermentation of Organic Residues to Beneficial Chemicals: A Review of Medium-Chain Fatty Acid Production. Processes 2020, 8 (12), 1571. 10.3390/pr8121571.

(17) Chen, W.-S.; Strik, D. P. B. T. B.; Buisman, C. J. N.; Kroeze, C. Production of Caproic Acid from Mixed Organic Waste: An Environmental Life Cycle Perspective. Environ. Sci. Technol. 2017, 51 (12), 7159–7168. 10.1021/acs.est.6b06220.

(18) Bazyar Lakeh, A. A.; Azizi, A.; Hosseini Koupaie, E.; Bekmuradov, V.; Hafez, H.; Elbeshbishy, E. A Comprehensive Study for Characteristics, Acidogenic Fermentation, and Anaerobic Digestion of Source Separated Organics. Journal of Cleaner Production 2019, 228, 73–85. 10.1016/j.jclepro.2019.04.223.

(19) Azizi, A.; Hosseini Koupaie, E.; Hafez, H.; Elbeshbishy, E. Improving Single- and Two-Stage Anaerobic Digestion of Source Separated Organics by Hydrothermal Pretreatment. Biochemical Engineering Journal 2019, 148, 77–86. 10.1016/j.bej.2019.05.001.

(20) Razavi, A. S.; Hosseini Koupaie, E.; Azizi, A.; Hafez, H.; Elbeshbishy, E. Hydrothermal Pretreatment of Source Separated Organics for Enhanced Solubilization and Biomethane Recovery. Bioresource Technology 2019, 274, 502–511. 10.1016/j.biortech.2018.12.024.

(21) Sukphun, P.; Sittijunda, S.; Reungsang, A. Volatile Fatty Acid Production from Organic Waste with the Emphasis on Membrane-Based Recovery. Fermentation 2021, 7 (3), 159. 10.3390/fermentation7030159.

(22) Hernandez, P. A.; Zhou, M.; Vassilev, I.; Freguia, S.; Zhang, Y.; Keller, J.; Ledezma, P.; Virdis, B. Selective Extraction of Medium-Chain Carboxylic Acids by Electrodialysis and Phase Separation. ACS Omega 2021, 6 (11), 7841–7850. 10.1021/acsomega.1c00397.

(23) Xu, J.; Bian, B.; Angenent, L. T.; Saikaly, P. E. Long-Term Continuous Extraction of Medium-Chain Carboxylates by Pertraction With Submerged Hollow-Fiber Membranes. Front Bioeng Biotechnol 2021, 9, 726946. 10.3389/fbioe.2021.726946.

(24) Kucek, L. A.; Nguyen, M.; Angenent, L. T. Conversion of L-Lactate into n-Caproate by a Continuously Fed Reactor Microbiome. Water Research 2016, 93, 163–171. 10.1016/j.watres.2016.02.018.

(25) Carvajal-Arroyo, J. M.; Andersen, S. J.; Ganigué, R.; Rozendal, R. A.; Angenent, L. T.; Rabaey, K. Production and Extraction of Medium Chain Carboxylic Acids at a Semi-Pilot Scale. Chemical Engineering Journal 2021, 416, 127886. 10.1016/j.cej.2020.127886.

(26) Agler, M. T.; Spirito, C. M.; Usack, J. G.; Werner, J. J.; Angenent, L. T. Chain Elongation with Reactor Microbiomes: Upgrading Dilute Ethanol to Medium-Chain Carboxylates. Energy Environ. Sci. 2012, 5 (8), 8189. 10.1039/c2ee22101b.

(27) Zhu, J.; Gutierrez, R. L.; Zhang, Y.; Sarker, N. R.; Dyussekenova, D.; Parmar, J.; Kim, B.-C.; Abbas, B.; Ren, K.; Lawson, C. E.; Werber, J. R. Selective and Solvent-Free Extraction of Medium-Chain Carboxylic Acids with Poly(Dimethylsiloxane) Membranes. ACS EST Eng. 2026, 6 (1), 460–472. 10.1021/acsestengg.5c00919.

(28) Xu, J.; Guzman, J. J. L.; Andersen, S. J.; Rabaey, K.; Angenent, L. T. In-Line and Selective Phase Separation of Medium-Chain Carboxylic Acids Using Membrane Electrolysis. Chem. Commun. 2015, 51 (31), 6847–6850. 10.1039/C5CC01897H.

(29) Xu, J.; Guzman, J. J. L.; Angenent, L. T. Direct Medium-Chain Carboxylic Acid Oil Separation from a Bioreactor by an Electrodialysis/Phase Separation Cell. Environ. Sci. Technol. 2021, 55 (1), 634–644. 10.1021/acs.est.0c04939.

(30) Norouzi, O.; Dutta, A. The Current Status and Future Potential of Biogas Production from Canada’s Organic Fraction Municipal Solid Waste. Energies 2022, 15 (2), 475. 10.3390/en15020475.

(31) 2540 SOLIDS. In Standard Methods For the Examination of Water and Wastewater. 10.2105/SMWW.2882.030.

(32) Klindworth, A.; Pruesse, E.; Schweer, T.; Peplies, J.; Quast, C.; Horn, M.; Glöckner, F. O. Evaluation of General 16S Ribosomal RNA Gene PCR Primers for Classical and Next-Generation Sequencing-Based Diversity Studies. Nucleic Acids Res 2013, 41 (1), e1. 10.1093/nar/gks808.

(33) Dueholm, M. K. D.; Andersen, K. S.; Korntved, A.-K. C.; Rudkjøbing, V.; Alves, M.; Bajón-Fernández, Y.; Batstone, D.; Butler, C.; Cruz, M. C.; Davidsson, Å.; Erijman, L.; Holliger, C.; Koch, K.; Kreuzinger, N.; Lee, C.; Lyberatos, G.; Mutnuri, S.; O’Flaherty, V.; Oleskowicz-Popiel, P.; Pokorna, D.; Rajal, V.; Recktenwald, M.; Rodríguez, J.; Saikaly, P. E.; Tooker, N.; Vierheilig, J.; De Vrieze, J.; Wurzbacher, C.; Nielsen, P. H. MiDAS 5: Global Diversity of Bacteria and Archaea in Anaerobic Digesters. Nat Commun 2024, 15 (1), 5361. 10.1038/s41467-024-49641-y.

(34) Lin, X.; Waring, K.; Ghezzi, H.; Tropini, C.; Tyson, J.; Ziels, R. M. High Accuracy Meets High Throughput for near Full-Length 16S Ribosomal RNA Amplicon Sequencing on the Nanopore Platform. PNAS Nexus 2024, 3 (10), pgae411. 10.1093/pnasnexus/pgae411.

(35) Bahram, M.; Anslan, S.; Hildebrand, F.; Bork, P.; Tedersoo, L. Newly Designed 16S rRNA Metabarcoding Primers Amplify Diverse and Novel Archaeal Taxa from the Environment. Environmental Microbiology Reports 2019, 11 (4), 487–494. 10.1111/1758-2229.12684.

(36) Scarborough, M. J.; Lawson, C. E.; Hamilton, J. J.; Donohue, T. J.; Noguera, D. R. Metatranscriptomic and Thermodynamic Insights into Medium-Chain Fatty Acid Production Using an Anaerobic Microbiome. mSystems 2018, 3 (6),. 10.1128/msystems.00221-18.

(37) Wang, Y.; Wu, J.; Lv, M.; Shao, Z.; Hungwe, M.; Wang, J.; Bai, X.; Xie, J.; Wang, Y.; Geng, W. Metabolism Characteristics of Lactic Acid Bacteria and the Expanding Applications in Food Industry. Front. Bioeng. Biotechnol. 2021, 9. 10.3389/fbioe.2021.612285.

(38) Chen, T.; Long, W.; Zhang, C.; Liu, S.; Zhao, L.; Hamaker, B. R. Fiber-Utilizing Capacity Varies in Prevotella-versus Bacteroides-Dominated Gut Microbiota. Sci Rep 2017, 7 (1), 2594. 10.1038/s41598-017-02995-4.

(39) Betancur-Murillo, C. L.; Aguilar-Marín, S. B.; Jovel, J. Prevotella: A Key Player in Ruminal Metabolism. Microorganisms 2022, 11 (1), 1. 10.3390/microorganisms11010001.

(40) Strachan, C. R.; Bowers, C. M.; Kim, B.-C.; Movsesijan, T.; Neubauer, V.; Mueller, A. J.; Yu, X. A.; Pereira, F. C.; Nagl, V.; Faas, J.; Wagner, M.; Zebeli, Q.; Weimer, P. J.; Candry, P.; Polz, M. F.; Lawson, C. E.; Selberherr, E. Distinct Lactate Utilization Strategies Drive Niche Differentiation between Two Co-Existing Megasphaera Species in the Rumen Microbiome. ISME J 2025, 19 (1), wraf147. 10.1093/ismejo/wraf147.

(41) Marounek, M.; Fliegrova, K.; Bartos, S. Metabolism and Some Characteristics of Ruminal Strains of Megasphaera Elsdenii. Appl Environ Microbiol 1989, 55 (6), 1570–1573. 10.1128/aem.55.6.1570-1573.1989.

(42) Kaminsky, R. A.; Reid, P. M.; Altermann, E.; Kenters, N.; Kelly, W. J.; Noel, S. J.; Attwood, G. T.; Janssen, P. H. Rumen Lachnospiraceae Isolate NK3A20 Exhibits Metabolic Flexibility in Response to Substrate and Coculture with a Methanogen. Applied and Environmental Microbiology 2023, 89 (10), e00634–23. 10.1128/aem.00634-23.

(43) Patel, G. B.; Sprott, G. D. Methanosaeta Concilii Gen. Nov., Sp. Nov. (“Methanothrix Concilii”) and Methanosaeta Thermoacetophila Nom. Rev., Comb. Nov.†. International Journal of Systematic and Evolutionary Microbiology 1990, 40 (1), 79–82. 10.1099/00207713-40-1-79.

(44) Ma, K.; Liu, X.; Dong, X. Methanosaeta Harundinacea Sp. Nov., a Novel Acetate-Scavenging Methanogen Isolated from a UASB Reactor. International Journal of Systematic and Evolutionary Microbiology 2006, 56 (1), 127–131. 10.1099/ijs.0.63887-0.

(45) Carr, S. A.; Schubotz, F.; Dunbar, R. B.; Mills, C. T.; Dias, R.; Summons, R. E.; Mandernack, K. W. Acetoclastic Methanosaeta Are Dominant Methanogens in Organic-Rich Antarctic Marine Sediments. ISME J 2018, 12 (2), 330–342. 10.1038/ismej.2017.150.

(46) Sauder, L. A.; Albertsen, M.; Engel, K.; Schwarz, J.; Nielsen, P. H.; Wagner, M.; Neufeld, J. D. Cultivation and Characterization of Candidatus Nitrosocosmicus Exaquare, an Ammonia-Oxidizing Archaeon from a Municipal Wastewater Treatment System. ISME J 2017, 11 (5), 1142–1157. 10.1038/ismej.2016.192.

(47) Liu, B.; Sträuber, H.; Saraiva, J.; Harms, H.; Silva, S. G.; Kasmanas, J. C.; Kleinsteuber, S.; Nunes da Rocha, U. Machine Learning-Assisted Identification of Bioindicators Predicts Medium-Chain Carboxylate Production Performance of an Anaerobic Mixed Culture. Microbiome 2022, 10 (1), 48. 10.1186/s40168-021-01219-2.

(48) Wallace, R. J.; McKain, N.; McEwan, N. R.; Miyagawa, E.; Chaudhary, L. C.; King, T. P.; Walker, N. D.; Apajalahti, J. H. A.; Newbold, C. J. Eubacterium Pyruvativorans Sp. Nov., a Novel Non-Saccharolytic Anaerobe from the Rumen That Ferments Pyruvate and Amino Acids, Forms Caproate and Utilizes Acetate and Propionate. International Journal of Systematic and Evolutionary Microbiology 2003, 53 (4), 965–970. 10.1099/ijs.0.02110-0.

(49) Seshadri, R.; Leahy, S. C.; Attwood, G. T.; Teh, K. H.; Lambie, S. C.; Cookson, A. L.; Eloe-Fadrosh, E. A.; Pavlopoulos, G. A.; Hadjithomas, M.; Varghese, N. J.; Paez-Espino, D.; Perry, R.; Henderson, G.; Creevey, C. J.; Terrapon, N.; Lapebie, P.; Drula, E.; Lombard, V.; Rubin, E.; Kyrpides, N. C.; Henrissat, B.; Woyke, T.; Ivanova, N. N.; Kelly, W. J. Cultivation and Sequencing of Rumen Microbiome Members from the Hungate1000 Collection. Nat Biotechnol 2018, 36 (4), 359–367. 10.1038/nbt.4110.

(50) Grootscholten, T. I. M.; Strik, D. P. B. T. B.; Steinbusch, K. J. J.; Buisman, C. J. N.; Hamelers, H. V. M. Two-Stage Medium Chain Fatty Acid (MCFA) Production from Municipal Solid Waste and Ethanol. Applied Energy 2014, 116, 223–229. 10.1016/j.apenergy.2013.11.061.

