## Supporting Information for "Long-term Production and Recovery of Medium-Chain Carboxylates from Source-Separated Organics"

### S1. Bioreactor Inoculum

At the Dufferin organics processing facility, the suspension buffer tank (SBT) is located between the grit removal system and the anaerobic digesters and controls the hydraulic flow of processed source-separated organics (SSO) pulp into the digesters (neither pH nor temperature is controlled). As a result, some degradation of the SSO pulp occurs before it enters the digesters. Due to the high concentrations of medium-chain carboxylic acids (MCCAs) observed in the SBT (Figure S1), this sample was chosen to be the bioreactor inoculum. Sieved SBT was mixed with sieved Dufferin SSO pulp in a 1:1 volumetric ratio as the start-up mixture in the bioreactor on Day 1 of operation. The characteristics of the inoculum (SBT sample) and SSO samples throughout the operational periods are provided in Table S1. Values are reported as mean ± standard deviation and represent temporal variations within each operating period.

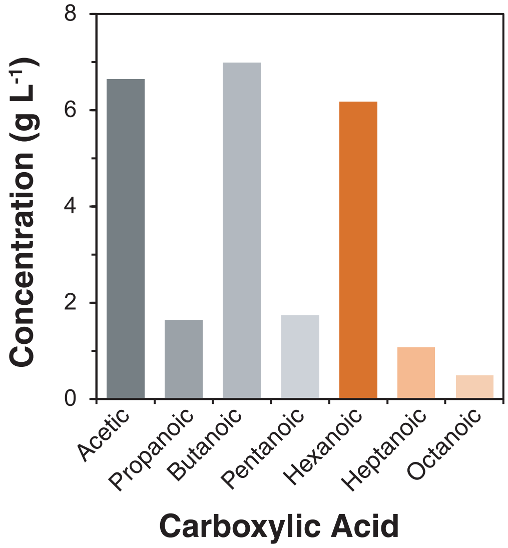


**Figure S1.** Concentration of C2–C8 carboxylic acids in a Suspension Buffer Tank sample from Dufferin organics processing facility. Concentrations of MCCAs are highlighted in shades of orange. This sample was part of the bioreactor inoculum on Day 1 of operation.

**Table S1.** Physical and chemical characteristics of the inoculum and feed SSO pulp samples from Dufferin and Disco Road[a,b]

| Parameter | Inoculum | Feed SSO Pulp | | | | | | | | |
|---|---|---|---|---|---|---|---|---|---|---|
| Operating Period | 1 | 1 | 1 | 2a | 2b | 3a | 3b | 4a | 4b | 5 |
| Day of Operation | 1 | 1 | 56–146 | 147–364 | 365–441 | 442–565 | 566–624 | 625–762 | 763–811 | 812–911 |
| SSO Feedstock | Dufferin | Dufferin | Dufferin | Dufferin | Dufferin | Disco Road | Disco Road | Disco Road | Disco Road | Disco Road |
| Outdoor Collection Temperature (°C) | 13.3 | 13.3 | 22.4 ± 2.7 | 3.6 ± 6.5 | 16.8 | 19.2 ± 2.3 | -1.4 ± 5.3 | 2.1 ± 10.8 | 21.4 ± 0.6 | 21.1 ± 0.6 |
| pH | 5.93 | 4.47 | N.D. | N.D. | N.D. | N.D. | 4.77 ± 0.22 | 4.71 ± 0.11 | 5.42 ± 0.10 | 5.61 ± 0.12 |
| TS (g $L^{-1}$) | 45.40 | 106.2 | 56.77 ± 10.65 | 55.52 ± 5.06 | 53.87 ± 4.32 | 36.25 ± 3.00 | 48.5 ± 1.49 | 45.70 ± 2.81 | 42.39 ± 2.81 | 35.21 ± 1.75 |
| VS (g $L^{-1}$) | 27.90 | 76.16 | 39.44 ± 7.84 | 39.58 ± 4.37 | 37.40 ± 4.53 | 23.26 ± 2.20 | 34.95 ± 0.76 | 33.43 ± 2.15 | 27.43 ± 2.30 | 21.97 ± 1.45 |
| TSS (g $L^{-1}$) | N.D. | N.D. | N.D. | 24.32 ± 3.29 | 22.09 ± 2.57 | 19.08 ± 3.18 | 22.53 ± 1.29 | 19.31 ± 1.62 | N.D. | 17.62 ± 0.82 |
| VSS (g $L^{-1}$) | N.D. | N.D. | N.D. | N.D. | 17.79 ± 1.83 | 15.59 ± 2.30 | 19.21 ± 1.22 | 17.60 ± 1.51 | N.D. | 14.92 ± 0.81 |
| Total COD (g COD $L^{-1}$) | N.D. | N.D. | 62.73 ± 12.26 | 72.92 ± 8.82 | 76.63 ± 4.53 | 58.36 ± 6.08 | 69.63 ± 4.43 | 65.41 ± 5.28 | 74.57 ± 4.47 | 69.11 ± 2.54 |
| Soluble COD (g COD $L^{-1}$) | N.D. | N.D. | 37.99 ± 7.57 | 38.73 ± 5.83 | 40.64 ± 2.60 | 32.76 ± 2.13 | 31.28 ± 1.04 | 32.48 ± 3.38 | 37.60 ± 1.31 | 39.44 ± 1.36 |
| Total Carbohydrates (g COD $L^{-1}$) | N.D. | N.D. | 10.94 ± 2.40 | 10.31 ± 1.12 | 5.53 ± 1.78 | 4.28 ± 1.12 | 7.09 ± 0.61 | 8.41 ± 0.79 | 5.07 ± 1.36 | 3.75 ± 0.78 |
| Soluble Carbohydrates (g COD $L^{-1}$) | N.D. | N.D. | 2.19 ± 0.51 | 2.42 ± 0.58 | 1.15 ± 0.10 | 0.75 ± 0.21 | 1.78 ± 0.37 | 2.06 ± 0.66 | 0.87 ± 0.29 | 0.55 ± 0.12 |
| Total Protein (g COD $L^{-1}$) | N.D. | 22.20 | 18.66 ± 2.81 | 18.50 ± 2.10 | 17.10 ± 1.16 | 16.35 ± 1.17 | 16.55 ± 0.30 | 14.67 ± 1.72 | 17.99 ± 0.22 | 18.87 ± 1.06 |
| Soluble Protein (g COD $L^{-1}$) | N.D. | 3.53 | 4.39 ± 0.49 | 3.78 ± 0.30 | 4.46 ± 0.14 | 3.77 ± 0.27 | 3.86 ± 0.38 | 3.84 ± 0.40 | 4.48 ± 0.59 | 4.22 ± 0.51 |
| SCCA (g COD $L^{-1}$) | 25.74 | 3.95 | 6.54 ± 1.70 | 6.11 ± 2.63 | 9.34 ± 1.66 | 16.73 ± 4.67 | 6.38 ± 0.96 | 5.12 ± 1.30 | 20.22 ± 4.29 | 26.14 ± 2.36 |

| | | | | | | | | | |
|---|---|---|---|---|---|---|---|---|---|
| Lactate (g COD $L^{-1}$) | 0.76 | 20.86 | 15.30 ± 4.46 | 16.77 ± 3.72 | 13.69 ± 0.90 | 4.45 ± 3.50 | 15.04 ± 0.87 | 15.39 ± 1.43 | 4.80 ± 2.29 | 0.64 ± 0.94 |
| Ethanol (g COD $L^{-1}$) | N.D. | 4.71 | 4.10 ± 1.66 | 4.19 ± 1.34 | 5.79 ± 0.91 | 3.77 ± 1.53 | 1.01 ± 0.31 | 1.28 ± 0.47 | 2.04 ± 0.30 | 2.26 ± 0.46 |

[a]Abbreviations: TS = total solids, VS = volatile solids, TSS = total suspended solids, VSS = volatile suspended solids, SCCA = short-chain carboxylic acids, N.D. = not determined. [b]Values are presented as mean ± standard deviation and represent temporal variations within each operating period; Day 1 values represent a single characterization of the inoculum and Dufferin SSO sample and are not period averages.

**S2. Sulfate Data**

The bioreactor pH was controlled at 5 by automated addition of 1 M sulfuric acid and/or 1 M potassium hydroxide (~5 mmol $H_2SO_4$ added $L^{-1}$ $day^{-1}$, Days 261–447). Sulfate concentrations were monitored during Days 261–447 to evaluate whether sulfate reduction occurred in the system. Sulfate concentrations were determined using an ion chromatograph (Dionex Integrion HPIC, Thermo Scientific, Waltham, MA, USA) equipped with a Dionex IonPac AS18 column (Thermo Scientific). Based on influent and effluent sulfate rates, no substantial sulfate reduction was observed during the monitored period (Figure S2). Influent rate was calculated as the sum of sulfate loading with SSO feed and acid dosing; effluent rate was calculated as sulfate concentration in the bioreactor divided by the SRT.

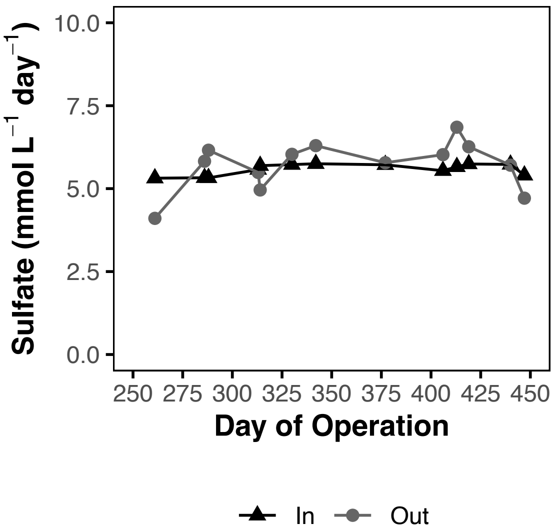


Figure S2. Sulfate influent rates (triangles; includes feeding rate with SSO and acid dosing with 1 M sulfuric acid) and effluent rates (circles; based on the measurements in the bioreactor) from Day 261–447.

### S3. In-line Extraction Preliminary Trials

The in-line extraction was integrated with the bioreactor during Days 557–563 (Trial 1) and Days 574–582 (Trial 2) as preliminary trials. During both trials, ultrafiltration (UF) membranes (1 module, 0.045 $m^2$ membrane surface area) submerged in the bioreactor were used for solids-liquid separation, and polydimethylsiloxane (PDMS) membranes (1 module, 0.1 $m^2$ membrane surface area) were used for MCCA extraction. An alkaline solution with a starting pH of 13 was used to drive the extraction during Trials 1 and 2 (manually controlled). During Trial 1, a vacuum was created on the UF permeate line (the pump was located after the PDMS membrane module), resulting in alkaline solution transfer to the bioreactor on Day 563 (pH of the bioreactor was maintained at 5). To restore the working volume to 3.4 L, UF permeate was wasted. During Trial 2, the pump was located before the PDMS membrane module. During both trials, MCCA extraction and PDMS efficiencies were low, 0.26 ± 0.12 g $g^{-1}$ and 0.22 ± 0.12 g $g^{-1}$, respectively, suggesting there was not enough PDMS membrane area to drive the extraction. Therefore, the PDMS membrane area was increased to 1.66 $m^2$ on Day 625, marking the start of continuous in-line extraction (Period 4a).

### S4. MCCA Oil Phase Separation

MCCAs were continuously extracted from the bioreactor in the alkaline solution controlled at a pH of 9 with automated dosing of 3 M NaOH during Period 4a (~5.4 mol NaOH added per mol MCCA extracted, Days 665–762). MCCA oil phase separation was performed twice (Trials A and B) using an alkaline solution concentrated with extracted carboxylic acids from Days 730–771. Phase separation included acidification of the alkaline bottle with 6 M hydrochloric acid (while mixing) to reach a final pH of 2, followed by settling and separation of the organic (oil) and aqueous phases. Each phase was characterized by chemical oxygen demand (COD) measurement and carboxylic acids quantification using GC-MS (Table S4-1).

In Trial A, 50 mL of an alkaline solution was acidified with 10 mL of acid (~5.5 mol HCl added per mol MCCA), resulting in 1 mL of phase-separated oil with an estimated mass of ~0.9 g, based on the measured oil density of 897.8 g $L^{-1}$. In Trial B, 800 mL of the same alkaline solution was acidified with 125 mL of acid, however, volumes of the resulting oil and aqueous phases were not measured. Therefore, because both trials originated from the same alkaline solution, the mass of carboxylic acids in each phase for Trial B was estimated using the volume proportions measured in Trial A, together with carboxylic acid concentrations measured in Trial B (Table S4-1). The distribution of MCCAs and short-chain carboxylic acids (SCCAs), relative to total carboxylic acids (TCA), was comparable between the two trials (Figure S4).

**Table S4-1.** Chemical and physical characteristics of alkaline bottle (pre-acidification), oil and aqueous layers (Trials A and B)

| Trial | Parameter | Unit | Alkaline Bottle | Oil | Aqueous |
|---|---|---|---|---|---|
| A | Volume | L | 0.05 | 0.001 | 0.059 |
| | Density | g $L^{-1}$ | 1044.6 | 897.8 | 1029.2 |
| | C2 | g $L^{-1}$ | 1.4 | 0.0 | 0.9 |
| | C3 | g $L^{-1}$ | 1.2 | 0.0 | 0.7 |
| | C4 | g $L^{-1}$ | 3.0 | 17.4 | 1.7 |
| | C5 | g $L^{-1}$ | 2.4 | 40.0 | 0.8 |
| | C6 | g $L^{-1}$ | 17.8 | 476.7 | 2.1 |
| | C7 | g $L^{-1}$ | 4.8 | 153.3 | 0.1 |
| | C8 | g $L^{-1}$ | 4.6 | 159.8 | 0.0 |
| | COD | g COD $L^{-1}$ | 76.7 | 2028.3 | 20.8 |
| B | Volume | L | 0.8 | N.D. | N.D. |
| | Density | g $L^{-1}$ | N.D. | N.D. | N.D. |
| | C2 | g $L^{-1}$ | 1.6 | 0.0 | 1.3 |
| | C3 | g $L^{-1}$ | 1.4 | 0.0 | 1.0 |
| | C4 | g $L^{-1}$ | 3.5 | 20.9 | 2.1 |
| | C5 | g $L^{-1}$ | 2.5 | 30.4 | 0.9 |
| | C6 | g $L^{-1}$ | 19.7 | 449.5 | 2.9 |
| | C7 | g $L^{-1}$ | 5.6 | 219.0 | 0.4 |
| | C8 | g $L^{-1}$ | 5.7 | 283.2 | 0.3 |
| | COD | g COD $L^{-1}$ | N.D. | 2255.0 | N.D. |

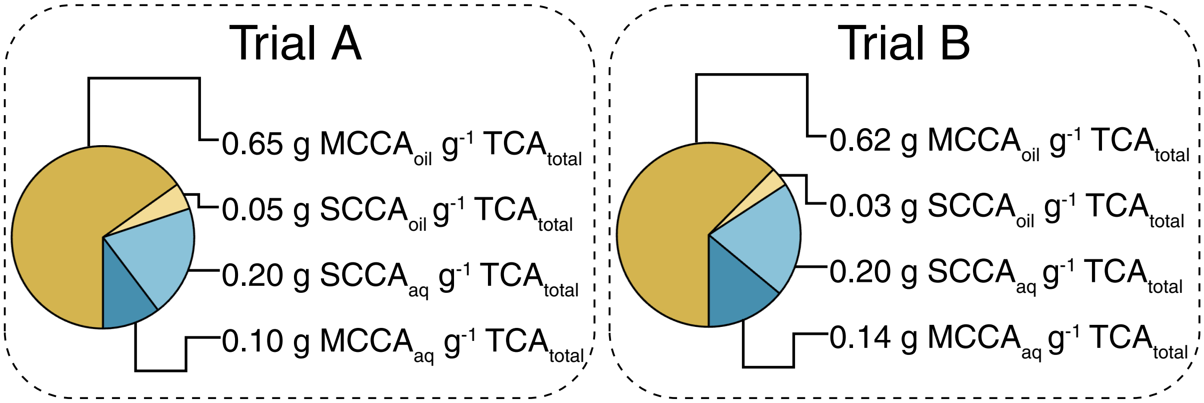


**Figure S4.** Fraction of MCCAs and short-chain carboxylic acids (SCCAs) per total carboxylic acids (TCA) post acidification in oil and aqueous layers (Trials A and B).

MCCAs recovered per total MCCA in the alkaline bottle in Trials A and B were 0.86 g $MCCA_{oil}$ $g^{-1}$ $MCCA_{total}$ and 0.82 g $MCCA_{oil}$ $g^{-1}$ $MCCA_{total}$, respectively. To estimate the MCCA concentration required to achieve 95% recovery, an iterative mass-balance calculation was performed using experimentally measured carboxylic acid distributions and densities from Trial A (Table S4-1) and previously reported solubilities of carboxylic acids (60 g $L^{-1}$ C4, 24 g $L^{-1}$ C5, 10.82 g $L^{-1}$ C6, 2.4 g $L^{-1}$ C7, 0.68 g $L^{-1}$ C8) [1–4]. An MCCA concentration of 200 g $L^{-1}$ in the alkaline bottle and a total volume of 1 L were assumed.

The concentration of total carboxylic acids (TCA) in the alkaline bottle was calculated using the measured distribution of 0.773 g MCCA $g^{-1}$ TCA (Trial A, Table S4-1):

$$[TCA_{alkaline}] = 200\ g\ MCCA\ L^{-1} \times \frac{g\ TCA}{0.773\ g\ MCCA} = 258.732\ g\ TCA\ L^{-1}$$

The mass of each carboxylic acid in the alkaline bottle was estimated using the measured carboxylic acid distribution in Trial A (Table S4-1). An example for C6 is shown below, and calculated values are tabulated in Table S4-2:

$$C6_{alkaline} = 258.732\ g\ TCA\ L^{-1} \times \frac{0.507\ g\ C6}{g\ TCA} \times 1\ L = \ 131.177\ g\ C6$$

The mass of each carboxylic acid in the aqueous phase was estimated assuming an aqueous phase volume of 0.79 L and constrained by reported solubility limits. If the solubility-limited mass was lower than the mass available in the alkaline bottle, the aqueous-phase mass was assumed to equal the solubility limit. An example for C6 is shown below:

$$C6_{solubility\ limit} = 10.82\ g\ C6\ L^{-1} \times 0.79\ L = 8.548\ g\ C6$$

$$C6_{solubility\ limit} < C6_{alkaline}$$

$$C6_{aqueous} = C6_{solubility\ limit}$$

Masses of the aqueous and oil phases were calculated using measured densities from Trial A (Table S4-1):

$$mass_{aqueous} = 0.79\ L\ \times 1029.2\ g\ L^{-1} = 813\ g$$

$$mass_{oil} = (1\ L - 0.79\ L) \times 897.8\ g\ L^{-1} = 189\ g$$

The mass of each carboxylic acid in the oil phase was estimated by the difference between the alkaline solution and aqueous phase. An example for C6 is shown below:

$$C6_{oil} = 131.177\ g\ C6 - 8.548\ g\ C6 = 122.629\ g\ C6$$

As a consistency check, the calculated total MCCA mass in the oil phase was compared with the calculated oil-phase mass, assuming the oil phase consisted only of MCCAs (Table S4-2). MCCA recovery was calculated as:

$$MCCA\ recovery = \frac{189\ g\ MCCA}{200\ g\ MCCA\ L^{-1} \times 1\ L} = 0.95$$

Based on this mass-balance approach, an MCCA concentration of approximately 200 g $L^{-1}$ in the alkaline bottle was estimated to achieve 95% MCCA recovery in the oil phase.

**Table S4-2.** Assumed and calculated values used to estimate the MCCA concentration required to achieve 95% recovery in the oil phase

| Parameter | Unit | Alkaline Bottle | Aqueous | Oil |
|---|---|---|---|---|
| Assumed Volume | L | 1 | 0.79 | 0.21 |
| Calculated Mass | g | - | 812.5 | 189[b] |
| C2 | g | 10.4 | 10.4 | 0 |
| C3 | g | 8.5 | 8.5 | 0 |
| C4 | g | 22.4 | 22.4 | 0 |
| C5 | g | 17.5 | 17.5 | 0 |
| C6 | g | 131.2 | 8.5 | 122.6 |
| C7 | g | 34.9 | 1.9 | 33 |
| C8 | g | 33.9 | 0.5 | 33.4 |
| MCCA | g | 200[a] | 11 | 189.0[b] |
| TCA | g | 258.8 | 69.8 | 189.0 |

[a]Assumed MCCA concentration for iterative calculations. [b]Iterations were performed until these values were equal.

**S5. Calculations of Performance Metrics**

Hydraulic retention time (HRT) was defined as the bioreactor working volume (L) divided by the feed flow rate (L day$^{-1}$). Solids retention time (SRT) was defined as the bioreactor working volume (L) divided by the effluent flow rate (L day$^{-1}$). The organic loading rate (OLR) (g $VS_{feed}$ L$^{-1}$ day$^{-1}$) was calculated as the volatile solids (VS) concentration in the SSO feed sample (g $VS_{feed}$ L$^{-1}$) divided by the HRT (days). In the same manner, feeding rates (g L$^{-1}$ day$^{-1}$) were calculated as the concentration of the respective compound in feed (g L$^{-1}$) divided by the HRT (day). Effluent rates (g L$^{-1}$ day$^{-1}$) were calculated as the concentration of a respective compound in the bioreactor (g L$^{-1}$) divided by the SRT (days). At a decoupled HRT and SRT, liquid waste rates (g L$^{-1}$ day$^{-1}$) were calculated as the concentration of the respective compound in the liquid waste (g L$^{-1}$) times liquid waste flow rate (L day$^{-1}$) divided by the bioreactor working volume (L). The reduction and production metrics (g g$^{-1}$) for lactate, carbohydrates, proteins and ammonium were calculated as the feeding rate minus the effluent rate minus the liquid waste rate (net produced/consumed), divided by the feeding rate.

MCCA production rate $r_b$ (g L$^{-1}$ day$^{-1}$) with in-line extraction (from Day 625) was calculated based on the mass balance approach (S1). MCCA yield (g MCCA g$^{-1}$ $VS_{feed}$) was calculated as MCCA production rate (g L$^{-1}$ day$^{-1}$) divided by the OLR (g VS L$^{-1}$ day$^{-1}$). MCCA extraction efficiency (g g$^{-1}$) was calculated as the MCCA extraction rate (S2) divided by the MCCA production rate (S1) times the bioreactor volume (L). PDMS efficiency was calculated as the difference of MCCA concentrations in the bioreactor and the recycle stream, divided by the bioreactor MCCA concentration.

$$r_b = \frac{r_e + r_s + r_l - r_f}{V_b} \tag{S1}$$

$$r_e = (C_b - C_r) \times Q_p \tag{S2}$$

$$r_s = C_b \times Q_s$$

$$r_l = C_r \times Q_l$$

$$r_f = C_f \times Q_f$$

Here,

$r_b$ = MCCA production rate (g L$^{-1}$ day$^{-1}$)

$r_e$ = MCCA extraction rate (g day$^{-1}$)

$r_s$ = MCCA effluent rate (g day$^{-1}$)

$r_l$ = MCCA liquid waste rate (g day$^{-1}$)

$r_f$ = MCCA feeding rate (g day$^{-1}$)

$V_b$ = bioreactor working volume (L)

$C_b$ = MCCA concentration in the bioreactor (g L$^{-1}$)

$C_r$ = MCCA concentration in the recycle stream after PDMS membranes (g L$^{-1}$)

$C_f$ = MCCA concentration in the feed (g L$^{-1}$)

$Q_p$ = permeate flow rate (L day$^{-1}$)

$Q_s$ = effluent flow rate (L day$^{-1}$)

$Q_l$ = liquid waste flow rate (L day$^{-1}$)

$Q_f$ = feed flow rate (L day$^{-1}$)

Volatile solids reduction (VSR) included MCCA production rate and was calculated as follows:

$$VSR = \frac{VS_f - VS_s - VS_l + r_b}{VS_f}$$

Here,

$r_b$ = MCCA production rate (g L$^{-1}$ day$^{-1}$) (S1)

$VS_f$ = VS feeding rate (g L$^{-1}$ day$^{-1}$)

$VS_s$ = VS effluent rate (g L$^{-1}$ day$^{-1}$)

$VS_l$ = VS liquid waste rate (g L$^{-1}$ day$^{-1}$)

**S6. Carboxylic Acids and Lactate Quantification**

Carboxylic acids and ethanol were quantified using an Agilent 8890 gas chromatograph (GC) equipped with tandem mass spectrometry (7000D Triple Quadrupole, Agilent, Santa Clara, CA, USA), autosampler (PAL RSI 120 Series 2, Agilent), DB-FatWax column (Agilent) with helium as a carrier gas. Samples for carboxylic acids analysis were diluted with HPLC-grade water and formic acid to a final concentration of 0.1M. The run time for carboxylic acids quantification was 16 minutes with the following temperature gradient: 80°C for 1 minute, 20°C min$^{-1}$ ramp until 200°C, 5°C min$^{-1}$ until 210°C, and 20°C min$^{-1}$ until 250°C, then held for 5 minutes. The dynamic multiple reaction monitoring (dMRM) method was used in mass spectrometry. Electron ionization was used to fragment ions of interest. Precursor and product ions for each compound were selected using Agilent software (Optimizer). Ethanol quantification included diluting samples in HPLC-grade water. Samples were heated at 95°C for 40 minutes prior to injection of the gas phase on GC. The run time was 12 minutes, with a 1.5-minute solvent delay and the following temperature gradient: 80°C for 1 minute, 20°C $min^{-1}$ ramp until 200°C, then held for 5 minutes. The scan method was used for mass spectrometry, with 29–200amu mass range and a scan time of 200ms.

Lactate was measured using a high-performance liquid chromatograph (UltiMate 3000 HPLC System, Thermo Scientific, Waltham, MA, USA) with an Aminex HPX-87H column (Bio-Rad, Hercules, CA, USA). Samples were diluted in HPLC-grade water prior to the analysis. Mobile phase used 5 mM sulfuric acid with a flow rate of 0.6 mL $min^{-1}$. The column temperature was set to 50°C. Ultraviolet detection at 214 nm was used for lactate identification and quantification.

#### S7. Ammonium Data

Ammonium concentrations were determined using an ion chromatograph (Dionex Integrion HPIC, Thermo Scientific) equipped with a Dionex IonPac CS19 column (Thermo Scientific) and a colorimetric quantification kit (AAT Bioquest, Pleasanton, CA, USA). Consistency between measurements obtained using the two methods was verified. (Table S7).

**Table S7.** Ammonium data from IC and colorimetric quantification assay

| Sample | Ammonium (g $L^{-1}$) - IC | Ammonium (g $L^{-1}$) - Assay |
|---|---|---|
| Feed (Day 419) | 1.67 | 1.74 |
| Bioreactor (Day 419) | 1.73 | 1.84 |
| Feed (Day 440) | 1.61 | 1.70 |
| Bioreactor (Day 440) | 1.67 | 1.77 |
| Feed (Day 911) | 1.54 | 1.69 |
| Bioreactor (Day 911) | 2.80 | 2.85 |

#### S8. Reactor Headspace Gas Composition

Bioreactor headspace composition ($CO_2$, $CH_4$, $O_2$, $N_2$) was analyzed weekly on GC (Hewlett Packard 5890, Agilent) with a thermal conductivity detector (TCD) and CTR I Concentric Packed Column (Alltech, Nicholasville, KY, USA) using helium carrier gas, while $H_2$ was verified on selected days using external GC-TCD (Clarus 500, Perkin Elmer, Shelton, CT, USA) with 60/80 Carboxen 1000 column (Supelco, Bellefonte, PA, USA) and argon as carrier gas (Table S8). $H_2$ was estimated based on the headspace balance, by subtracting the sum of ($CO_2$, $CH_4$, $O_2$, $N_2$) from

100%, assuming atmospheric pressure inside the bioreactor (Table S8), since gas produced in the system was vented to the pressure bottle filled with water.

**Table S8.** Hydrogen estimated from the bioreactor headspace balance and verified measurements

| Day of Operation | $H_2$ % (estimated) | $H_2$ % (measured externally) |
|---|---|---|
| 208 | 37.75 | 34.98 |
| 210 | 37.20 | 31.42 |
| 310 | 0.21 | 6.39 |
| 324 | 30.75 | 15.17 |
| 337 | 29.84 | 33.13 |
| 373 | 20.83 | 29.37 |
| 462 | 0.00 | 0.74 |

On days when oxygen was introduced to the bioreactor, nitrogen gas was used to sparge the system. Additionally, once in-line extraction was implemented (Period 4a onwards), gas bags pre-filled with nitrogen were used to maintain atmospheric pressure in the system (attached to alkaline bottle and dosing bottles with acid/base). Therefore, on these days, gas headspace composition was normalized, assuming $N_2$ displaced $CO_2$ and $CH_4$ produced in the bioreactor (Figure S8). $N_2$ was recalculated based on the $O_2$ measurements, assuming the ratio of $O_2/N_2$ of 0.27 (air composition). Then, once methane production was detected, CO2 and CH4 were recalculated assuming equal displacement by nitrogen. On days when methane was not produced, $CO_2$ was recalculated assuming introduced $N_2$ displaced CO2.

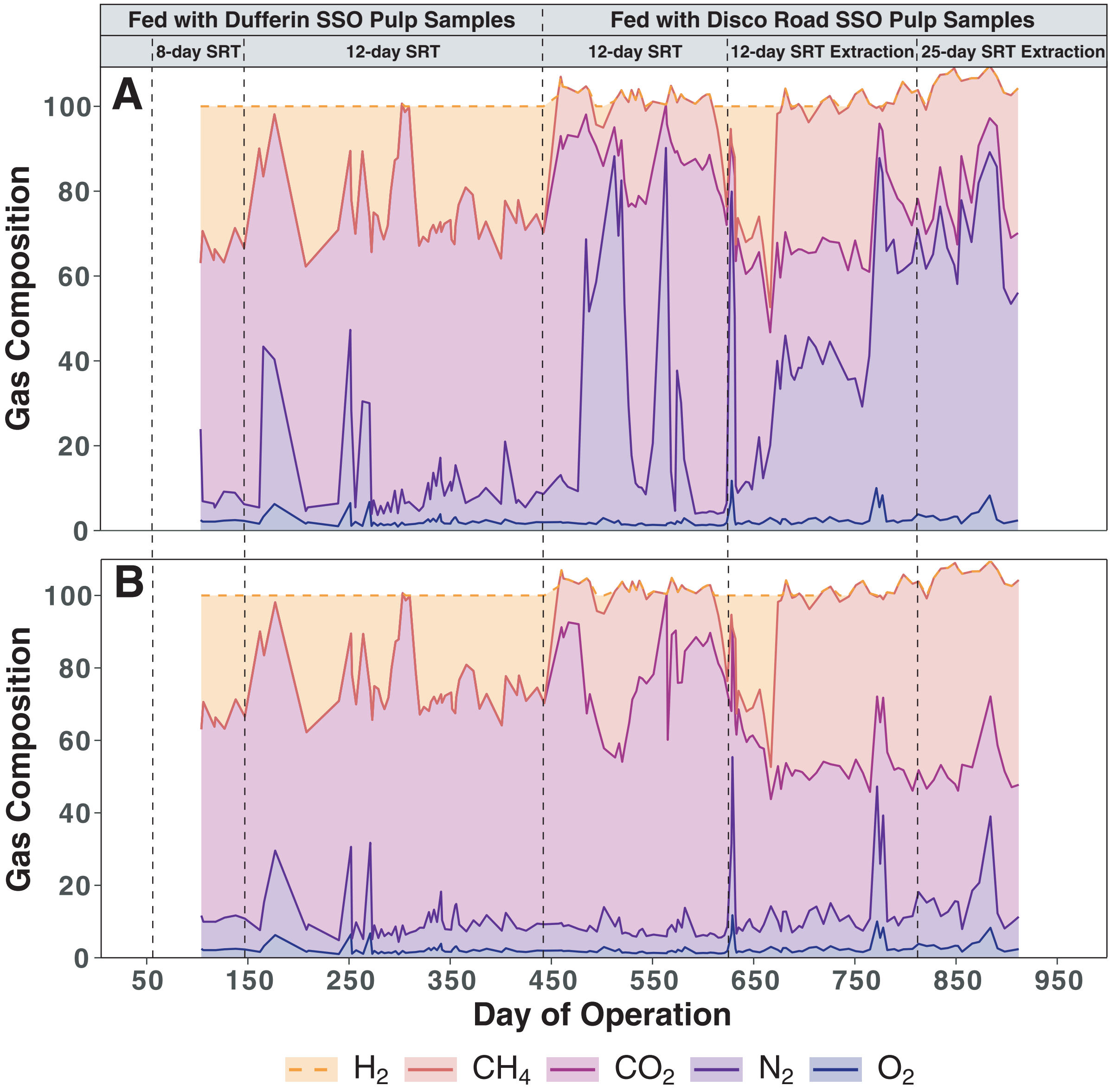


**Figure S8.** Bioreactor headspace composition as % gas: (A) nitrogen is not normalized, and (B) normalized for nitrogen introduced during bioreactor sparging and during in-line extraction operation.

### S9. Bioinformatics and Statistical Analyses of 16S rRNA Gene Sequencing Data

POD5 files were processed using Dorado (v1.3.0) and associated super-accurate basecalling model (dna_r10.4.1_e8.2_400bps_sup@v5.2.0) before being demultiplexed according to barcodes. The resulting reads were filtered to retain only sequences between 1.2 and 1.8 kbp for bacterial read sets and 1.0 to 1.2 kbp for archaeal read sets. Reads were subsequently quality controlled with NanoPack2's chopper (v0.12.0)[5] to remove reads with average quality scores less than Q15. Quality controlled reads were then taxonomically profiled at the genus level using Kraken2 (v2.1.4)[6] against SILVA (SSU v138).[7] To provide greater taxonomic resolution at the species level for select samples, we also analyzed bacterial read sets with Emu (v3.5.2)[8] using a custom database containing 16S rRNA reference sequences derived from the most recent release of GTDB (r226).[9]

Statistical analyses of microbial community data were performed in R (v.4.3.1) using the vegan (v. 2.7.1)[10] package. Alpha diversity was assessed using the Shannon diversity index. Bray–Curtis dissimilarities were calculated from genus-level relative abundance data and used for nonmetric multidimensional scaling (NMDS) ordination to compare microbial community composition of SSO feedstock and bioreactor samples across the eight operating periods. Differences in microbial community composition between groups were assessed using permutational multivariate analysis of variance (PERMANOVA) based on Bray–Curtis dissimilarities, while homogeneity of multivariate dispersion was evaluated using betadisper, as implemented in the vegan package.

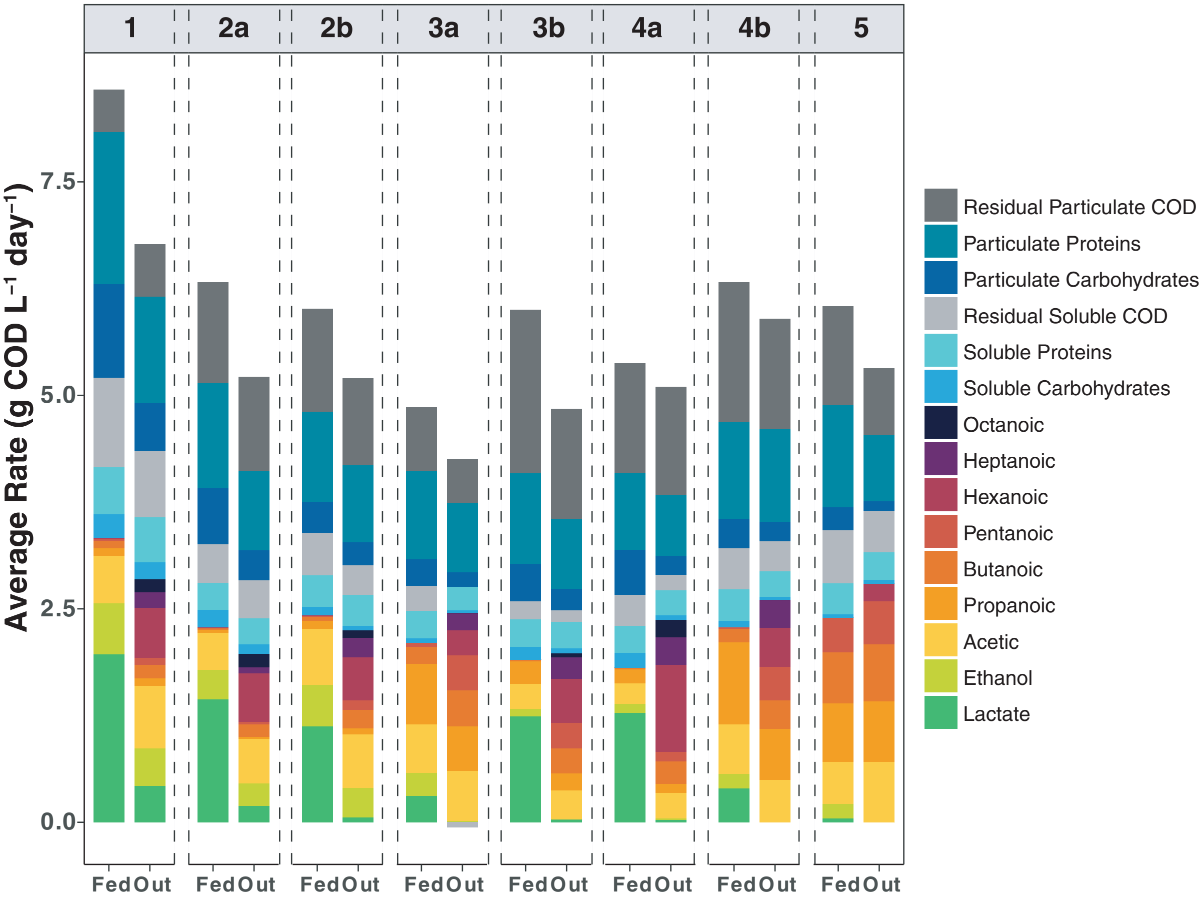


**Figure S10.** COD mass balance based on the feeding (fed) and effluent (out) rates across eight operating periods. Total bars represent total COD.

### S11. Operation at a 25-day SRT and 12-day HRT

The AnMBR operated at a decoupled SRT of 25 days and HRT of 12 days from Days 812–911 (Period 5, with extraction, warm Disco pulp samples). The SRT was increased to 25 days to evaluate whether longer biomass retention could improve the hydrolysis of the remaining organic matter present in the warm Disco pulp, characterized by reduced lactate availability and higher SCCA concentrations. Production rate of C2 was the highest across total carboxylic acids, reaching $0.21 \pm 0.10$ g $L^{-1}$ $day^{-1}$, and C3 production was first observed during this period. Production rate of C6 was the lowest at a 25-day SRT, with no production of C7 and C8. SCCA production rate was the highest, reaching $0.31 \pm 0.14$ g $L^{-1}$ $day^{-1}$ in Period 5, while MCCA production was the lowest ($0.09 \pm 0.03$ g $L^{-1}$ $day^{-1}$).

Although lactate was fully degraded in the feedstock before it was introduced to the bioreactor, there were still available carbon sources, including proteins and carbohydrates. Carbohydrate reduction (0.48 ± 0.08 g $g^{-1}$) was comparable to Periods 3a and 4b (warm Disco pulp samples), while the reduction of protein increased to 0.3 ± 0.07 g $g^{-1}$ and was the highest across all periods (Figure S11). Production of ammonia also increased to 0.49 ± 0.15 g $g^{-1}$ in this period. As proteins are hydrolyzed, they release amino acids, which are potential alternative electron donors that could be used for chain elongation. Since amino acids were not measured in this study, increased ammonia production provided indirect evidence of increased protein degradation. These results suggest that the longer SRT may have enhanced degradation of residual organic matter in the SSO, as reflected by increased protein reduction and ammonia production. However, increased hydrolysis was not accompanied by increased MCCA production (Figure S11).

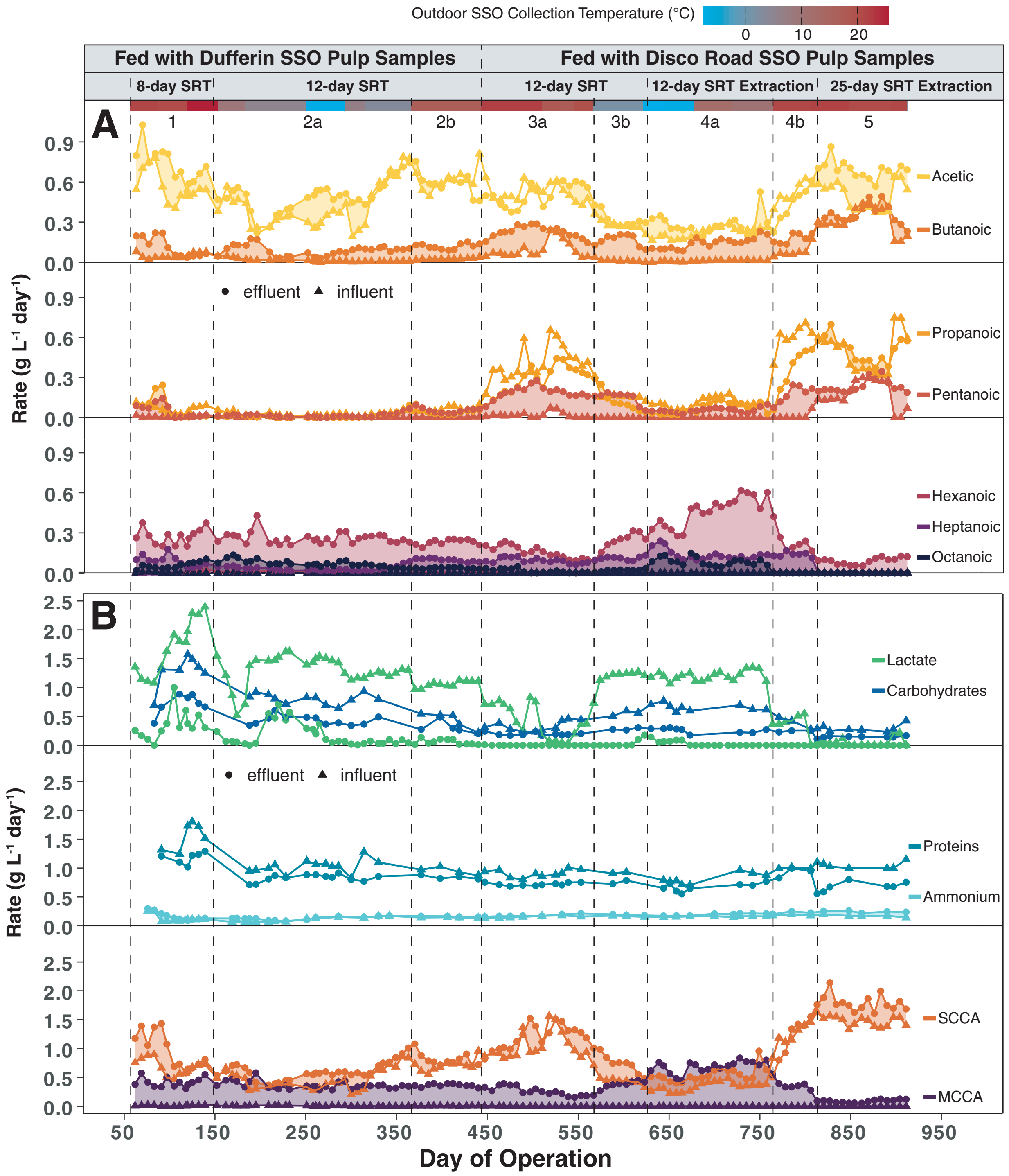


**Figure S11.** Bioreactor performance over operating periods (dashed vertical lines): (A) feeding (triangles) and effluent (circles) rates of carboxylic acids, with filled regions representing production rates, (B) feeding (triangles) and effluent (circles) rates of lactate, carbohydrates, proteins, ammonia, SCCAs and MCCAs. Carboxylic acids and lactate sample points are 5-day averages, while carbohydrates, proteins and ammonia are not averaged as characterized not daily. Effluent rates from Period 4a include extraction rates.

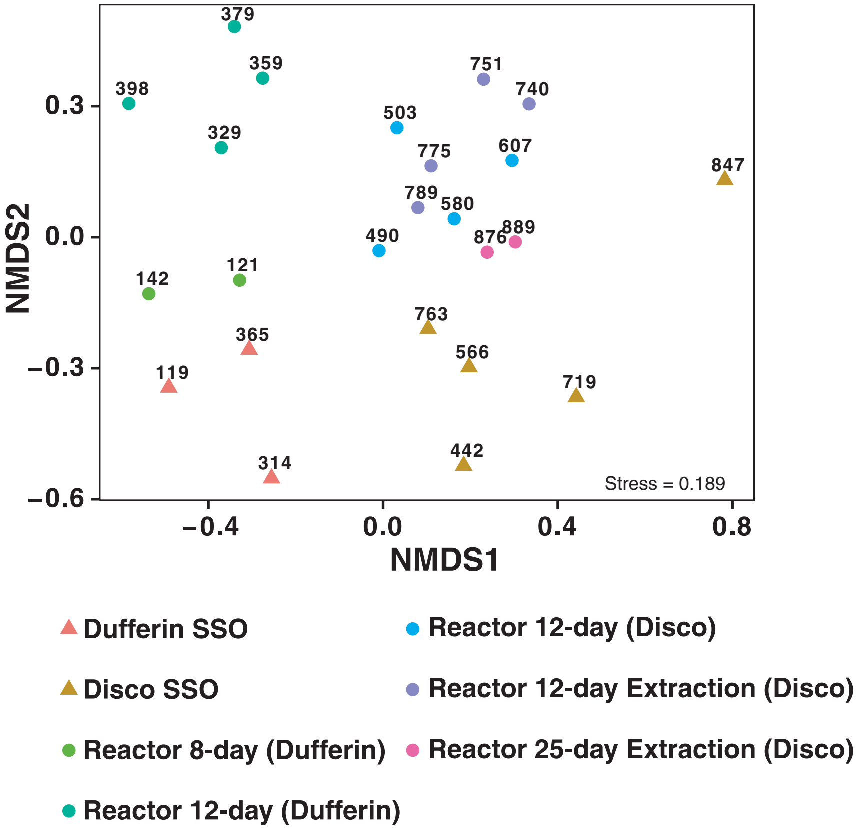


**Figure S12.** NMDS ordination analysis based on Bray–Curtis dissimilarities using genus-level 16S rRNA gene sequencing data in the SSO feedstock and bioreactor samples across eight operating periods. Numbers correspond to the Day of Operation.

**Table S13.** PERMANOVA and multivariate dispersion analyses of microbial community composition across feedstock and bioreactor sample categories

| Data sets | Category | PERMANOVA $R^2$ | PERMANOVA F | PERMANOVA p | Betadisper p |
|---|---|---|---|---|---|
| Feed and Bioreactor | Feed versus Bioreactor | 0.182 | 4.885 | 0.001 | 0.993 |
| Feed | Facility | 0.528 | 6.714 | 0.017 | 0.330 |
| Bioreactor | Facility | 0.526 | 15.547 | 0.002 | 0.412 |
| Bioreactor | Operational Mode (SRT, Extraction) | 0.460 | 3.403 | 0.003 | <0.001[a] |

[a]Significant betadisper results indicate unequal multivariate dispersion among groups

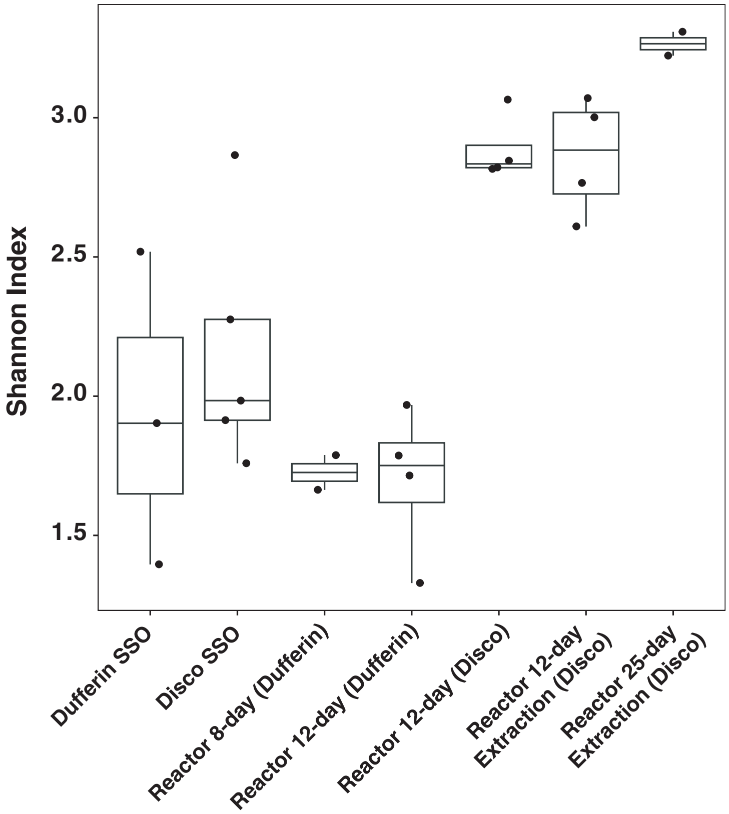


**Figure S14.** Alpha diversity (Shannon index) of SSO feedstock and bioreactor samples based on genus-level 16S rRNA gene sequencing data.

**Table S15.** Estimated theoretical C6 production rates versus measured values

| **Operating Period** | **1** | **2a** | **2b** | **3a** | **3b** | **4a** | **4b** | **5** |
|---|---|---|---|---|---|---|---|---|
| Days of Operation | 56–146 | 147–364 | 365–441 | 442–565 | 566–624 | 625–762 | 763–811 | 812–911 |
| SRT (days) | 8 | 12 | 12 | 12 | 12 | 12 | 12 | 25 |
| HRT (days) | 8 | 12 | 12 | 12 | 12 | 12 | 12 | 12 |
| In-line Extraction (+/-) | - | - | - | - | - | + | + | + |
| SSO Feedstock | Dufferin | Dufferin | Dufferin | Disco Road | Disco Road | Disco Road | Disco Road | Disco Road |
| Outdoor Collection Temperature (°C) | 22.4 ± 2.7 | 3.6 ± 6.5 | 16.8 | 19.2 ± 2.3 | -1.4 ± 5.3 | 2.1 ± 10.8 | 21.4 ± 0.6 | 21.1 ± 0.6 |
| Days of Pseudo Steady-State Operation | 79–146 (67 days) | 182–364 (182 days) | 365–441 (76 days) | 484–565 (81 days) | 566–624 (58 days) | 625–762 (137 days) | 765–811 (46 days) | 847–911 (64 days) |
| Measured Lactate Consumption Rate (g COD $L^{-1}$ $day^{-1}$) | 1.54 ± 0.43 | 1.25 ± 0.19 | 1.07 ± 0.10 | 0.31 ± 0.29 | 1.21 ± 0.18 | 1.25 ± 0.11 | 0.4 ± 0.21 | 0.04 ± 0.09 |
| Measured C6 Production Rate (g COD $L^{-1}$ $day^{-1}$) | 0.56 ± 0.12 | 0.56 ± 0.1 | 0.5 ± 0.05 | 0.29 ± 0.07 | 0.51 ± 0.16 | 1.02 ± 0.24 | 0.46 ± 0.09 | 0.21 ± 0.06 |
| Estimated C6 Production Rate[a] (g COD $L^{-1}$ $day^{-1}$) | 1.37 ± 0.38 | 1.11 ± 0.17 | 0.95 ± 0.09 | 0.28 ± 0.26 | 1.08 ± 0.16 | 1.12 ± 0.1 | 0.36 ± 0.19 | 0.04 ± 0.08 |

[a]Estimated based on 3 mol lactate to 1 mol C6 stoichiometry.[11]

### References


(1) Angenent, L. T.; Richter, H.; Buckel, W.; Spirito, C. M.; Steinbusch, K. J. J.; Plugge, C. M.; Strik, D. P. B. T. B.; Grootscholten, T. I. M.; Buisman, C. J. N.; Hamelers, H. V. M. Chain Elongation with Reactor Microbiomes: Open-Culture Biotechnology To Produce Biochemicals. *Environ. Sci. Technol.* **2016**, *50* (6), 2796–2810. https://doi.org/10.1021/acs.est.5b04847.

(2) O'Neil, M. J.; Co, M. &. *The Merck Index : An Encyclopedia of Chemicals, Drugs, and Biologicals*, 14th ed.; Merck: Whitehouse Station, N.J, 2006.

(3) Hemphill, L.; Swanson, W. S. Proceedings of the 18th Industrial Waste Conference; Engineering Bulletin; Purdue University: Lafayette, IN, 1964; Vol. 18, pp 204–217.

(4) Yalkowsky, S. H.; He, Y. *Handbook of Aqueous Solubility Data*; CRC Press: Boca Raton, FL, 2003.

(5) De Coster, W.; Rademakers, R. NanoPack2: Population-Scale Evaluation of Long-Read Sequencing Data. *Bioinformatics* **2023**, *39* (5), btad311. https://doi.org/10.1093/bioinformatics/btad311.

(6) Wood, D. E.; Lu, J.; Langmead, B. Improved Metagenomic Analysis with Kraken 2. *Genome Biol.* **2019**, *20*, 257. https://doi.org/10.1186/s13059-019-1891-0.

(7) Chuvochina, M.; Gerken, J.; Frentrup, M.; Sandikci, Y.; Goldmann, R.; Freese, H. M.; Göker, M.; Sikorski, J.; Yarza, P.; Quast, C.; Peplies, J.; Glöckner, F. O.; Reimer, L. C. SILVA in 2026: A Global Core Biodata Resource for rRNA within the DSMZ Digital Diversity. *Nucleic Acids Res.* **2026**, *54* (D1), D334–D341. https://doi.org/10.1093/nar/gkaf1247.

(8) Curry, K. D.; Wang, Q.; Nute, M. G.; Tyshaieva, A.; Reeves, E.; Soriano, S.; Wu, Q.; Graeber, E.; Finzer, P.; Mendling, W.; Savidge, T.; Villapol, S.; Dilthey, A.; Treangen, T. J. Emu: Species-Level Microbial Community Profiling of Full-Length 16S rRNA Oxford Nanopore Sequencing Data. *Nat. Methods* **2022**, *19* (7), 845–853. https://doi.org/10.1038/s41592-022-01520-4.

(9) Parks, D. H.; Chuvochina, M.; Rinke, C.; Mussig, A. J.; Chaumeil, P.-A.; Hugenholtz, P. GTDB: An Ongoing Census of Bacterial and Archaeal Diversity through a Phylogenetically Consistent, Rank Normalized and Complete Genome-Based Taxonomy. *Nucleic Acids Res.* **2022**, *50* (D1), D785–D794. https://doi.org/10.1093/nar/gkab776.

(10) Oksanen, J.; Simpson, G. L.; Blanchet, F. G.; Kindt, R.; Legendre, P.; Minchin, P. R.; O'Hara, R. B.; Solymos, P.; Stevens, M. H. H.; Szoecs, E.; Wagner, H.; Barbour, M.; Bedward, M.; Bolker, B.; Borcard, D.; Borman, T.; Carvalho, G.; Chirico, M.; Caceres, M. D.; Durand, S.; Evangelista, H. B. A.; FitzJohn, R.; Friendly, M.; Furneaux, B.; Hannigan, G.; Hill, M. O.; Lahti, L.; Martino, C.; McGlinn, D.; Ouellette, M.-H.; Cunha, E. R.; Smith, T.; Stier, A.; Braak, C. J. F. T.; Weedon, J. *Vegan: Community Ecology Package*; 2025.

(11) Scarborough, M. J.; Lawson, C. E.; Hamilton, J. J.; Donohue, T. J.; Noguera, D. R. Metatranscriptomic and Thermodynamic Insights into Medium-Chain Fatty Acid Production Using an Anaerobic Microbiome. *mSystems* **2018**, *3* (6), 10.1128/msystems.00221-18. https://doi.org/10.1128/msystems.00221-18.
